# Heterodimers of functionally divergent ARF-GEF paralogues prevented by self-interacting dimerisation domain

**DOI:** 10.1101/2021.09.14.460214

**Authors:** Sabine Brumm, Manoj K. Singh, Hauke Beckmann, Sandra Richter, Kerstin Huhn, Tim Kucera, Sarah Baumann, Choy Kriechbaum, Hanno Wolters, Shinobu Takada, Gerd Jürgens

**Author notes:** These authors contributed equally (Sabine Brumm, Manoj K. Singh).

## Abstract

Functionally divergent paralogs of homomeric proteins do not form potentially deleterious heteromers, which requires distinction between self and non-self (Hochberg et al., 2018; Marchant et al, 2019; Marsh and Teichmann, 2015). In Arabidopsis, two ARF guanine-nucleotide exchange factors (ARF-GEFs) related to mammalian GBF1, named GNOM and GNL1, can mediate coatomer complex (COPI)-coated vesicle formation in retrograde Golgi-endoplasmic reticulum (ER) traffic (Geldner et al., 2003; Richter et al., 2007; Teh and Moore, 2007). Unlike GNL1, however, GNOM is also required for polar recycling of endocytosed auxin efflux regulator PIN1 from endosomes to the plasma membrane. Here we show that these paralogues form homodimers constitutively but no heterodimers. We also address why and how GNOM and GNL1 might be kept separate. These paralogues share a common domain organisation and each N-terminal dimerisation (DCB) domain can interact with the complementary fragment (ΔDCB) of its own and the other protein. However, unlike self-interacting DCB^GNOM^ (Grebe et al., 2000; Anders et al., 2008), DCB^GNL1^ did not interact with itself nor DCB^GNOM^. DCB^GNOM^ removal or replacement with DCB^GNL1^, but not disruption of cysteine bridges that stabilise DCB-DCB interaction, resulted in GNOM-GNL1 heterodimers which impaired developmental processes such as lateral root formation. We propose precocious self-interaction of the DCB^GNOM^ domain as a mechanism to preclude formation of fitness-reducing GNOM-GNL1 heterodimers.

## Introduction

ARF guanine-nucleotide exchange factors (ARF-GEFs) promote the formation of transport vesicles on endomembranes by catalysing the GDP-GTP exchange of small ARF GTPases through their SEC7 domain (Mossessova et al., 2003; Renault et al., 2003; reviewed in Casanova, 2007; Anders and Jürgens, 2008). Plant genomes only encode large ARF-GEFs, which are evolutionarily conserved among eukaryotes and have a distinct domain organisation (Cox et al., 2004; Mouratou et al., 2005; Anders and Jürgens, 2008; Bui et al., 2009; Pipaliya et al., 2019). The centrally located catalytic SEC7 domain is flanked by a Homology Upstream of SEC7 (HUS) domain and three or four Homology Downstream of SEC7 (HDS1-3 or HDS1-4) domains (Mouratou et al., 2005; Anders and Jürgens, 2008). In addition, there is an N-terminal dimerisation and cyclophilin-binding (DCB) domain which in several ARF-GEFs has been shown to interact with itself and at least one other domain of the same ARF-GEF (Grebe et al., 2000; Ramaen et al., 2007; Anders et al., 2008). In Arabidopsis, interactions involving the DCB domain are required for membrane association of ARF-GEF GNOM, functional complementation of mutant GNOM proteins and formation of functional GNOM dimers mediating coordinated activation of ARF1 GTPases (Anders et al., 2008; Brumm et al., 2020).

In Arabidopsis, there are three paralogues related to human GBF1. Like GBF1, GNOM and GNOM-LIKE 1 (GNL1) each can mediate COPI traffic from the Golgi stacks to the ER whereas GNOM but not GNL1 is also required for polar recycling of auxin efflux carrier PIN1 from endosomes to the basal plasma membrane (Steinmann et al., 1999; Geldner et al., 2003; Richter et al., 2007; Teh and Moore, 2007). The third paralogue GNL2 essentially behaves like GNOM but is specifically expressed and required in haploid pollen development (Richter et al., 2011). GNOM and GNL1 co-exist in virtually all tissues and yet only GNOM performs the task of polar recycling of PIN1. This is remarkable because GNOM and GNL1 are closely related by sequence, with 60% of their respective 1451 and 1443 amino acid residues being identical. Their functional divergence suggests that GNOM and GNL1 are kept apart within the cell. Here we demonstrate that GNL1, like GNOM, forms homodimers but does not form heterodimers with GNOM which, when engineered, impair development and then address how GNOM and GNL1 might be kept separate.

## Results

Co-immunoprecipitation revealed interaction of differently tagged GNL1 proteins in transgenic Arabidopsis seedling extract, much like that of GNOM (Fig. 1A,B). In contrast, no GNL1-GNOM heteromers were detected (Fig. 1B). ARF-GEFs are cytosolic and associate with endomembranes for activation of their ARF substrates (Steinmann et al., 1999; Anders et al., 2008). Cell fractionation followed by co-immunoprecipitation demonstrated that like GNOM (Brumm et al., 2020), GNL1 proteins exist as homomers both in the cytosol and on membranes, indicating that these proteins form homomers constitutively rather than only in the context of membrane association (Fig. 1C).

**Figure 1.**
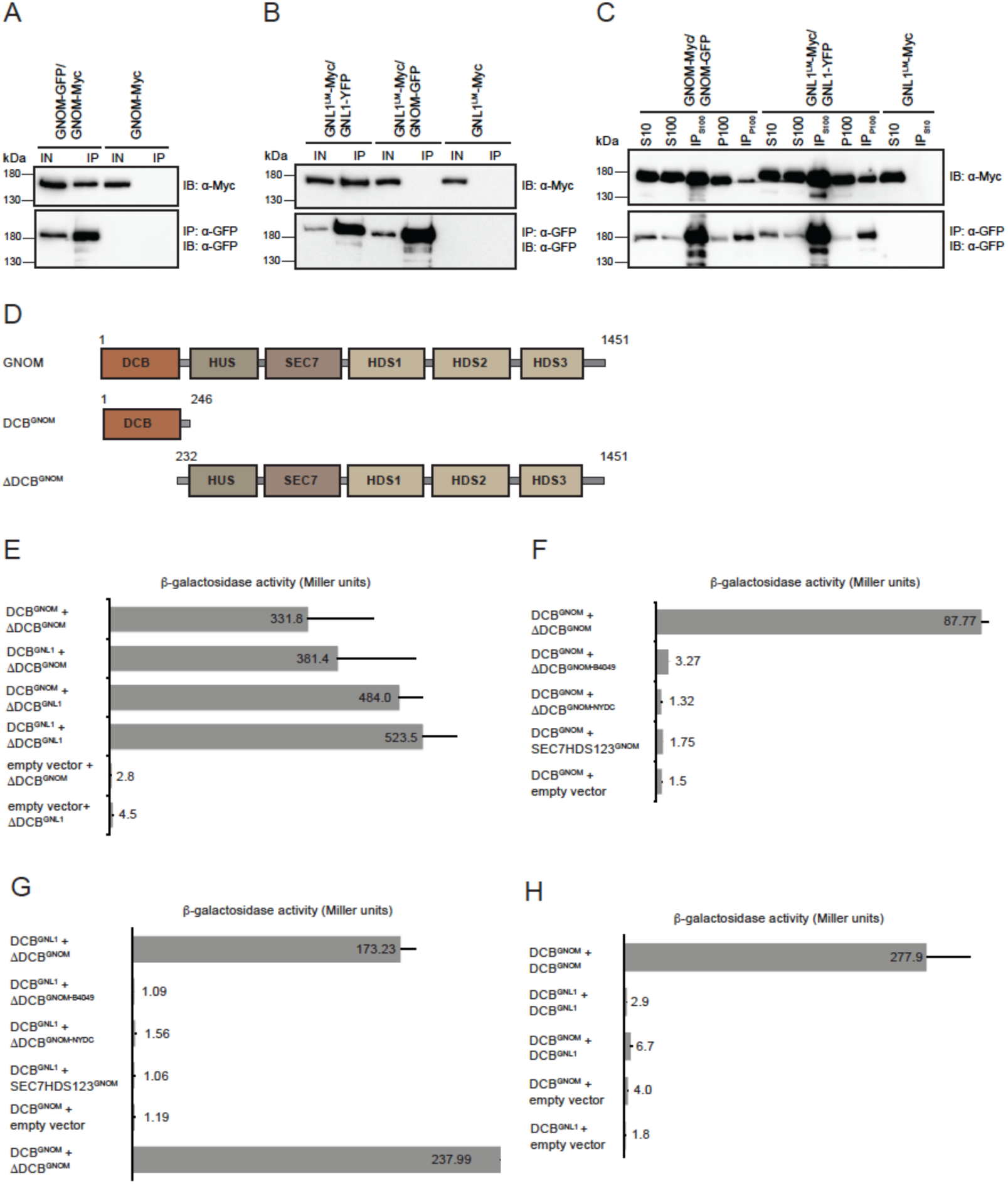
Paralogous ARF-GEFs GNOM and GNL1 – no heteromer formation but domain interaction. **(A-C)** In-planta co-immunoprecipitation interaction assays of full-length proteins. IN, input; IP, immunoprecipitate. Proteins were separated by SDS-PAGE and probed with specific antisera (IB; right); protein sizes in kDa (left). (**A**) Interaction of GNOM-Myc with GNOM-HA. GNOM-Myc, negative control. (**B**) Interaction of GNL1^LM^-Myc with GNL1-YFP but no GNOM-GNL1 interaction. GNL1^LM^-Myc, negative control. GNL1^LM^, engineered BFA-sensitive variant of GNL1 (Richter et al., 2007). (**C**) Cell fractionation and co-IP of differently tagged GNL1 from Arabidopsis seedlings. S10, S100, P100, supernatants and pellet from centrifugation at 10,000 x *g* and 100,000 x *g*. GNOM-GFP x GNOM-Myc, positive control; GNL1^LM^-Myc, negative control. **(D-H)** Quantitative yeast two-hybrid interaction assays of DCB domain (**D**) Diagram of domain organisation of ARF-GEFs GNOM and GNL1. The DCB^GNOM^ domain spans aa1-246, the complementary ΔDCB^GNOM^ fragment (comprising domains HUS, SEC7, HDS1, HDS2 and HDS3) spans aa232-1451. (**E**) Both DCB^GNOM^ and DCB^GNL1^ interacted with ΔDCB^GNL1^ and ΔDCB^GNOM^. (**F, G**) Interaction of (**F**) DCB^GNOM^ and (**G**) DCB^GNL1^ with wild-type ΔDCB^GNOM^. Both DCB domains failed to interact with ΔDCB^GNOM^ variants bearing HUS box (ΔDCB^GNOM-NYDC^) or G579R mutation (ΔDCB^GNOM-B4049^) or with a ΔDCB^GNOM^ fragment lacking the HUS domain (SEC7HDS123^GNOM^). (**H**) DCB^GNL1^ did not interact with itself, unlike DCB^GNOM^, nor with DCB^GNOM^. **Figure 1 – figure supplement 1.** DCB-DCB interaction assays of GNOM-GNL1 chimeric DCB domains

The domain organisation is identical between GNOM and GNL1. In GNOM, the N-terminal DCB domain interacts with the complementary ΔDCB fragment to mediate membrane association (Anders et al., 2008; see Fig. 1D). Using the yeast two-hybrid assay, we also detected DCB-ΔDCB interaction in GNL1, which was very similar to that in GNOM (Fig. 1D-G). Moreover, each DCB domain interacted not only with the ΔDCB fragment of its own protein but also with that of the paralogue (Fig. 1E). The two DCB domains also behaved identically in their interaction with truncated or mutated ΔDCB fragments of GNOM (Fig. 1F-G). These results suggest that the two proteins have the potential to form GNOM-GNL1 heteromers via DCB-ΔDCB interaction, which however, appears to be prevented by some unknown mechanism(s) and for unknown reason(s) in planta. Nonetheless, there was one specific difference between the two DCB domains. Only DCB^GNOM^ interacted with itself whereas DCB^GNL1^ did not interact with itself nor with DCB^GNOM^ (Fig. 1H). Thus, although both GNOM and GNL1 form homomers, GNL1 homomerisation relies on DCB-ΔDCB interaction only whereas GNOM homomerisation can also be mediated by DCB-DCB interaction.

Gel electrophoresis under non-reducing conditions revealed a distinct higher band for both GNOM and GNL1, consistent with the occurrence of homodimers (Fig. 2A). Exposure to 5 mM or more dithiothreitol (DTT) shifted the GNL1 band to the monomer size, further supporting the idea that cysteine bridges might be involved in stabilising the dimers (Fig. 2B). The DCB domain of GNOM has 7 cysteine residues (Fig. 2C). To assess their significance, we generated different sets of C-to-S substitutions, yielding GNOM^7CS^, GNOM^4CS^, GNOM^3CS^ and GNOM^1CS^, and tested the mutant DCB^GNOM^ variants in yeast two-hybrid experiments for their ability to interact with themselves, DCB^GNOM^ and ΔDCB^GNOM^ (Fig. 2C-E). All these mutant DCB^GNOM^ variants failed to interact with themselves and with the wild-type form of DCB^GNOM^ (Fig. 2D). Regarding the interaction with ΔDCB^GNOM^, only DCB^GNOM-3CS^ and the complementary substitution variant DCB^GNOM-1CS^ interacted whereas the other variants DCB^GNOM-4CS^ and DCB^GNOM-7CS^ failed to do so (Fig. 2E). Thus, the C-to-S substitutions impaired the interaction capability of the DCB^GNOM^ domain, with the DCB-DCB interaction apparently being more sensitive than the DCB-ΔDCB interaction. We then generated transgenic lines expressing full-length GNOM variants with C-to-S substitutions (Figure 2 – figure supplement 1).

**Figure 2.**
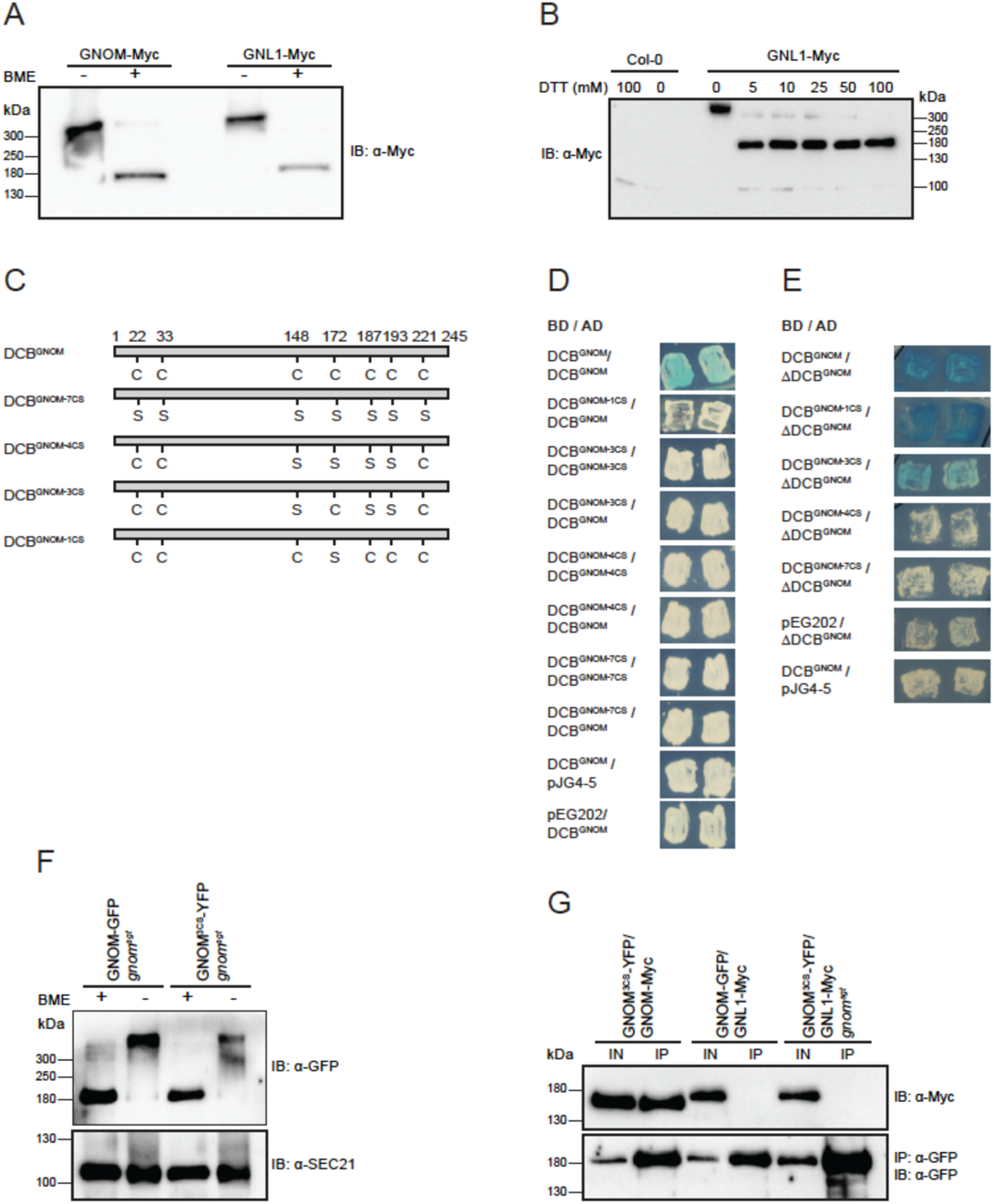
Interaction behaviour and functionality of C-to-S substitution mutants. (**A-B**) Redox-dependent GNOM and GNL1 dimer detection (**A**) Apparent dimers of GNOM and GNL1 detected in Western blots under non-reducing conditions (BME, ß-mercaptoethanol). (**B**) Band shift to monomer size in 5 mM or more dithiothreitol (DTT), suggesting involvement of cysteine bridges in stabilising the dimers. (**C-G**) Interaction behaviour of GNOM with C-to-S substitutions (GNOM^CS^) (**C**) Positions of C residues and their C-to-S substitutions indicated in DCB domain of wild-type and 3CS, 4CS, 7CS and 1CS mutant GNOM proteins. (**D-E**) Yeast two-hybrid interaction assays. (**D**) None of the mutant DCB^GNOM^ domains interacted with itself or with wild-type DCB^GNOM^ domain. (**E**) DCB^GNOM-3CS^ and DCB^GNOM-1CS^ interacted with the ΔDCB^GNOM^ fragment, like wild-type DCB^GNOM^ and in contrast to the other two C>S substitution mutants. BD, DNA-binding domain; AD, activation domain. (**F**) GNOM^3CS^ formed homodimers detectable under non-reducing conditions, although the dimer-representing bands appeared abnormal. (**G**) Co-immunoprecipitation analysis of GNOM^3CS^-YFP and GNL1-Myc from transgenic Arabidopsis seedling extract. **Figure 2 – figure supplement 1**. Expression of transgenes in *gnom^sgt^* background **Figure 2 – figure supplement 2**. Rescue of *gnom^sgt^* mutant plants with C-to-S substitution variants of GNOM **Figure 2 – figure supplement 3**. Subcellular localisation of GNOM^CS^ mutant proteins **Figure 2 – figure supplement 4**. Interaction behaviour of GNOM^4CS^ **Figure 2 – figure supplement 5**. Complementation of *gnom^sgt^ gnl1* double knockout mutant with GNOM^3CS^

Unexpectedly, full-length GNOM^4CS^ and GNOM^7CS^ mutant proteins rescued Arabidopsis plants lacking the endogenous *GNOM* gene (Figure 2 – figure supplement 2). In addition, GNOM^4CS^ associated with endosomal membranes in the absence of endogenous GNOM (Figure 2 – figure supplement 3), although there was no DCB^GNOM-4CS^-ΔDCB^GNOM^ interaction as required for membrane association (Fig. 2E). GNOM^4CS^ was also unable to interact with itself but did interact with GNOM and with GNOM^3CS^, indicating that the ΔDCB fragment of GNOM^4CS^ was able to interact with DCB domains other than its own (Figure 2 – figure supplement 4). In contrast, GNOM^3CS^ was not only able to interact with GNOM^4CS^ (Figure 2 – figure supplement 4) but also appeared to be functional on its own since it rescued not only the lethal *gnom^sgt^* mutant, a 37-kb deletion of *GNOM* and flanking genes (Brumm et al., 2020), but also the gametophytically lethal *gnom^sgt^ gnl1* double mutant lacking both paralogues (Figure 2 – figure supplement 2 and 5; Suppl. Table 1). However, the *gnom^sgt^ gnl1* double mutant rescued by GNOM^3CS^ showed a strong pollen transmission defect that revealed not only severe growth retardation but also impairment of both GNL1 and GNOM functions in pollen development (Suppl. Table 1). Interestingly, GNOM^3CS^ formed homodimers detectable under non-reducing conditions, although the dimer-representing bands appeared abnormal (Fig. 2F). Like GNOM, GNOM^3CS^ did not interact with GNL1, but interacted with GNOM (Fig. 2G). This suggests that the initial self-recognition of DCB^GNOM^ might not be affected by the 3CS substitution, although no stable DCB-DCB interaction of GNOM^3CS^ was detected in the yeast two-hybrid assay (see Fig. 2E). It is also noteworthy that all but one of the C residues of DCB^GNOM^ are conserved in DCB^GNL1^. This suggests a more general role of the C-C bridges in stabilising the structure of the DCB domain, which might support its ability to interact with another DCB domain and/or a ΔDCB fragment. In contrast to the C-to-S substitution variants with their critical residues in the C-terminal half of the DCB domain, chimeric DCB domains comprising complementary parts from GNOM and GNL1 revealed that the N-terminal aa1-144 determined the DCB-DCB interaction behaviour according to its origin (Figure 1 – figure supplement 1). In conclusion, the cysteine bridges of the DCB domain appear to have rather general roles in stabilising the interaction ability of DCB^GNOM^ and DCB^GNL1^, and in this way promote the functionality of both GNOM and GNL1.

Next we addressed whether DCB-DCB interaction is required for GNOM function and/or plays a role in preventing GNOM-GNL1 heterodimer formation by expressing, from the *GNOM* cis-regulatory region, a Myc-tagged chimeric variant that had DCB^GNL1^ in place of DCB^GNOM^ (designated DCB^GNL1^:ΔDCB^GNOM^-Myc) (Figure 2 – figure supplement 1). The chimera rescued both the *gnom^sgt^* deletion and the *gnom^sgt^ gnl1* double mutant (Figure 3 – figure supplement 1). However, the rescued *gnom^sgt^ gnl1* plants were strongly reduced in size and pollen development was impaired (Figure 3 – figure supplement 1; Suppl. Table 1). In the *gnom^sgt^* mutant expressing DCB^GNL1^:ΔDCB^GNOM^, lateral root development involving GNOM-dependent polar recycling of PIN1 (Fig. 3A) and GNOM-dependent root gravitropism appeared not to be affected (Fig 3B*, top row*). These results suggested that DCB-DCB interaction is not essential for GNOM function.

**Figure 3.**
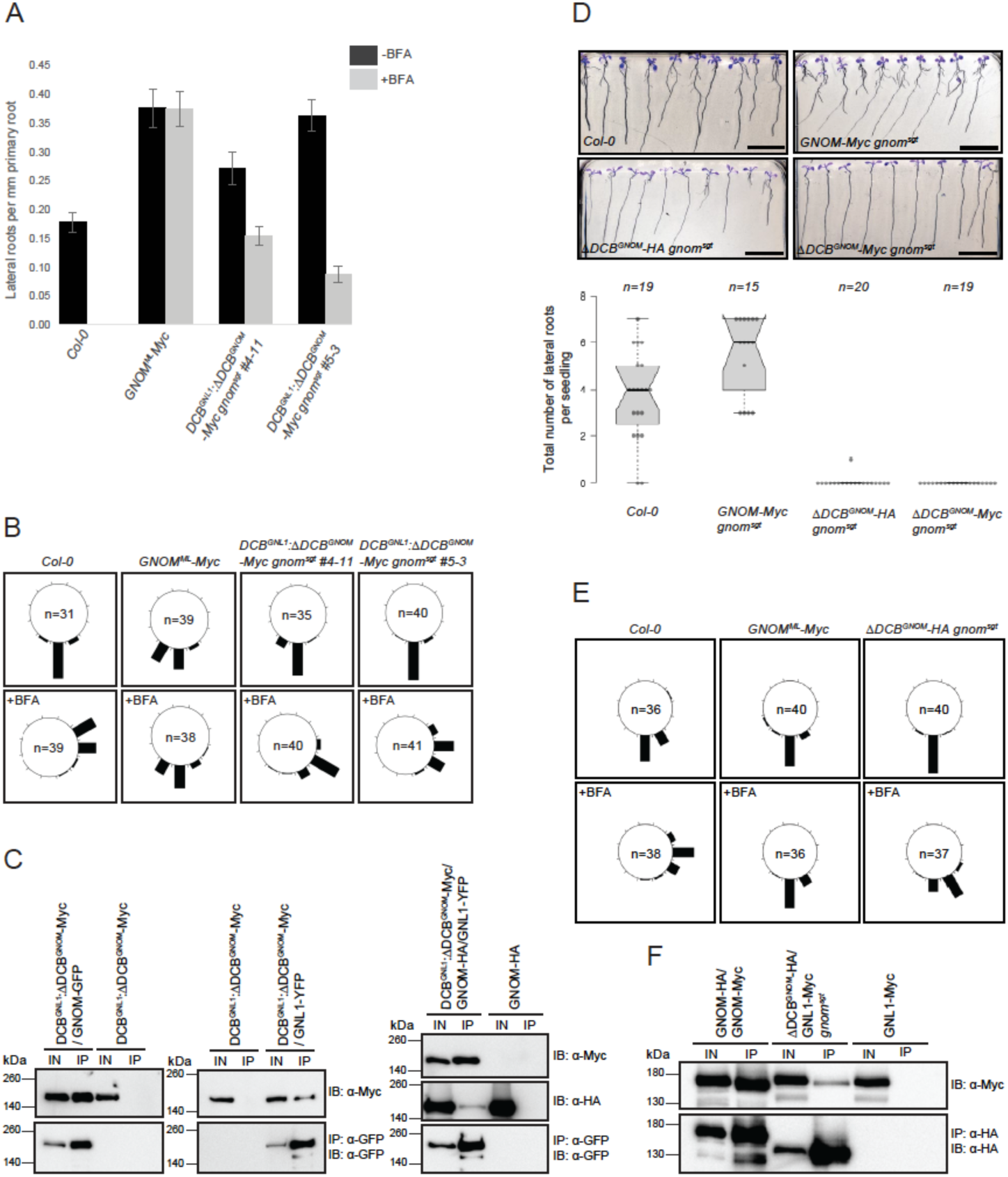
Developmental phenotypes and GNOM-GNL1 interaction in *gnom^sgt^* deletion mutants rescued by DCB^GNL1^:ΔDCB^GNOM^ chimeric protein or ΔDCB^GNOM^ fragment. **(A-C)** DCB^GNL1^:ΔDCB^GNOM^ protein. (**A**) Lateral root development and (**B**) root gravitropism normal in the absence of BFA and partially resistant to BFA due to interaction with BFA-resistant GNL1. Controls: Col-0, wild-type; GNOM^ML^-Myc, BFA-resistant GNOM. 10µM BFA. (**C**) Interaction of DCB^GNL1^:ΔDCB^GNOM^-Myc chimeric protein with both GNOM-GFP (*left*) and GNL1-YFP (*middle*). Heterotrimer (*right*) with DCB^GNL1^:ΔDCB^GNOM^-Myc acting as a bridge between GNOM-HA and GNL1-YFP. (**D-F**) HA-tagged or Myc-tagged ΔDCB^GNOM^ protein. (**D**) No lateral root development. Controls: Col-0, wild-type; GNOM^ML^-Myc, BFA-resistant GNOM. (**E**) Root gravitropism normal in the absence of BFA and nearly fully resistant to BFA due to interaction with BFA-resistant GNL1. Controls: Col-0, wild-type; GNOM^ML^-Myc, BFA-resistant GNOM. 10µM BFA. (**F**) GNOM without DCB domain (ΔDCB^GNOM^-HA) interacting with GNL1-Myc in *gnom^sgt^* homozygous background. (**C, F**) Protein extracts of Arabidopsis seedlings expressing differently tagged proteins were subjected to co-immunoprecipitation analysis. Total extracts (IN) and immunoprecipitates (IP) were separated by SDS-PAGE and probed with specific antisera (IB) indicated on the right; protein sizes are given in kDa on the left. **Figure 3 – figure supplement 1**. Postembryonic phenotypes of *gnom^sgt^* deletion mutant and *gnom^sgt^ gnl1* double mutant rescued by expression of chimeric DCB^GNL1^:ΔDCB^GNOM^ or ΔDCB^GNOM^ protein **Figure 3 – figure supplement 2**. NAA-induced lateral root initiation in *gnom^sgt^* seedlings rescued by ΔDCB^GNOM^

Then we subjected the chimera to co-immunoprecipitation analysis. The Myc-tagged DCB^GNL1^:ΔDCB^GNOM^ chimera interacted with both GNOM and GNL1 presumably because the lack of DCB-DCB interaction allowed for interaction of the chimera with the two endogenous paralogous ARF-GEFs (Fig. 3C). Moreover, GNOM was also co-immunoprecipitated with GNL1 in the presence of the chimera which thus appears to act as a bridging protein, enabling the formation of ARF-GEF heterotrimers (Fig. 3C, compare with Fig. 1B). The interaction of the chimera with GNL1 became functionally relevant when the seedlings were exposed to the fungal inhibitor brefeldin A (BFA). BFA inhibits the GDP-GTP exchange activity of GNOM whereas GNL1 is a BFA-resistant ARF-GEF (Geldner et al., 2003; Richter et al., 2007). Both lateral root development and root gravitropism were partially resistant to BFA in *gnom* mutant seedlings rescued by DCB^GNL1^:ΔDCB^GNOM^ in comparison to the BFA-sensitive wild-type control, consistent with the formation of DCB^GNL1^:ΔDCB^GNOM^-GNL1 heterodimers (Fig. 3A; Fig. 3B, *bottom row*). Assuming that GNL1 confers BFA resistance to the heterodimer, partial BFA resistance suggests that only some DCB^GNL1^:ΔDCB^GNOM^-GNL1 heterodimers are formed in addition to BFA-sensitive DCB^GNL1^:ΔDCB^GNOM^ homodimers. In conclusion, DCB-DCB interaction is not essential for GNOM function but appears to be involved in preventing the formation of GNOM-GNL1 heterodimers.

The occurrence of DCB^GNL1^:ΔDCB^GNOM^-GNL1 heterodimers made us consider the possibility of full-length GNL1 interacting with the GNOM fragment lacking the DCB domain (ΔDCB^GNOM^), which was expressed from the *GNOM* cis-regulatory region in the *gnom^sgt^* background (Figure 2 – figure supplement 1). Indeed, ΔDCB^GNOM^ was able to rescue the *gnom^sgt^* deletion mutant, which can be attributed to its interaction with GNL1 since DCB-ΔDCB interaction is necessary for membrane association and function of GNOM (Anders et al., 2008; Fig. 3F; Figure 3 – figure supplement 1). This interpretation was supported by the observation that root gravitropism was normal and almost fully resistant to BFA in *gnom^sgt^* mutant seedlings rescued by ΔDCB^GNOM^, very much like in engineered BFA-resistant GNOM and in contrast to the BFA-sensitive wild-type control (Fig. 3E, *bottom row*). This result indicates that the GNOM activity required for root gravitropism was entirely provided by ΔDCB^GNOM^-GNL1 heterodimers. Consistent with this, ΔDCB^GNOM^ did not rescue the *gnom^sgt^ gnl1* double mutant (Suppl. Table 2A). Similarly, ΔDCB^GNOM^ bearing the *B4049* mutation failed to rescue the *gnom^sgt^* deletion mutant (Suppl. Table 2B), which can be attributed to the failure of DCB^GNL1^ to interact with ΔDCB^GNOM-B4049^ (see Fig. 1G). In addition, unlike ΔDCB^GNOM^, ΔDCB^GNOM-B4049^ does not interact with full-length GNOM since the *B4049* mutation (G_579_R substitution) abolishes the DCB-ΔDCB interaction, and *B4049* consequently interferes with membrane-association of GNOM (Anders et al., 2008). Thus, in contrast to the chimeric protein DCB^GNL1^:ΔDCB^GNOM^ which was active on its own, the *gnom^sgt^* rescue of ΔDCB^GNOM^ required the interaction with the DCB domain of full-length GNL1. This interaction provided membrane association competence, thus revealing the endosomal targeting potential of ΔDCB^GNOM^. Importantly, although *gnom^sgt^* deletion plants were rescued by the *ΔDCB^GNOM^* transgene, they showed specific abnormalities. Lateral root formation was completely abolished in *ΔDCB^GNOM^* transgenic *gnom^sgt^* mutants in comparison to wildtype controls (Fig. 3D). Treatment of seedlings with the auxin analogue NAA over night or for 2 days promotes lateral root formation from pericycle cells in wildtype. In contrast, *gnom^sgt^* mutants rescued by ΔDCB^GNOM^ often displayed strong proliferation of pericycle cells, which frequently resulted in multiple lateral root primordia; however, only some of these primordia were almost shaped like wild-type primordia (Figure 3 – figure supplement 2) and no primordia developed into lateral roots (see Fig. 3D). These results suggest that lateral root development might be particularly sensitive to GNL1-ΔDCB^GNOM^ heterodimer formation since both GNOM-dependent endosomal recycling and GNL1-mediated secretion are simultaneously required during lateral root formation.

Because membrane association requires DCB-ΔDCB interaction (Anders et al., 2008), rescue of the *gnom^sgt^* deletion mutant by the ΔDCB^GNOM^ fragment would imply subcellular relocation of GNL1 from Golgi stacks to the endosomal membranes where GNOM mediates polar recycling of PIN1 to the basal plasma membrane (Fig. 4). Using ARF1 as a marker for the endosomal BFA compartment, we detected GNL1 in the surrounding Golgi stacks, which is its normal location (Fig. 4A-D). In the presence of ΔDCB^GNOM^, however, GNL1 co-localised with ARF1 very much like GNOM (Fig. 4E-H, compare with Fig. 4I-L). The relocation of GNL1 caused by ΔDCB^GNOM^ was also detectable using the Golgi marker γCOP as a reference which in addition, indicated that COPI recruitment was still functional, presumably due to the formation of GNL1-GNL1 homodimers (Fig. 4M-P, compare with Fig. 4Q-T). These observations suggest that by DCB-ΔDCB interaction with full-length GNL1, the ΔDCB^GNOM^ fragment gains the ability to associate with membranes and directs the heterodimer to endosomal membranes, thus providing GNOM activity. However, this rescue of GNOM-dependent recycling by GNL1 appears to be contingent upon the demand for GNL1-mediated secretion as seen for example in lateral root development (see Fig. 3D).

**Figure 4.**
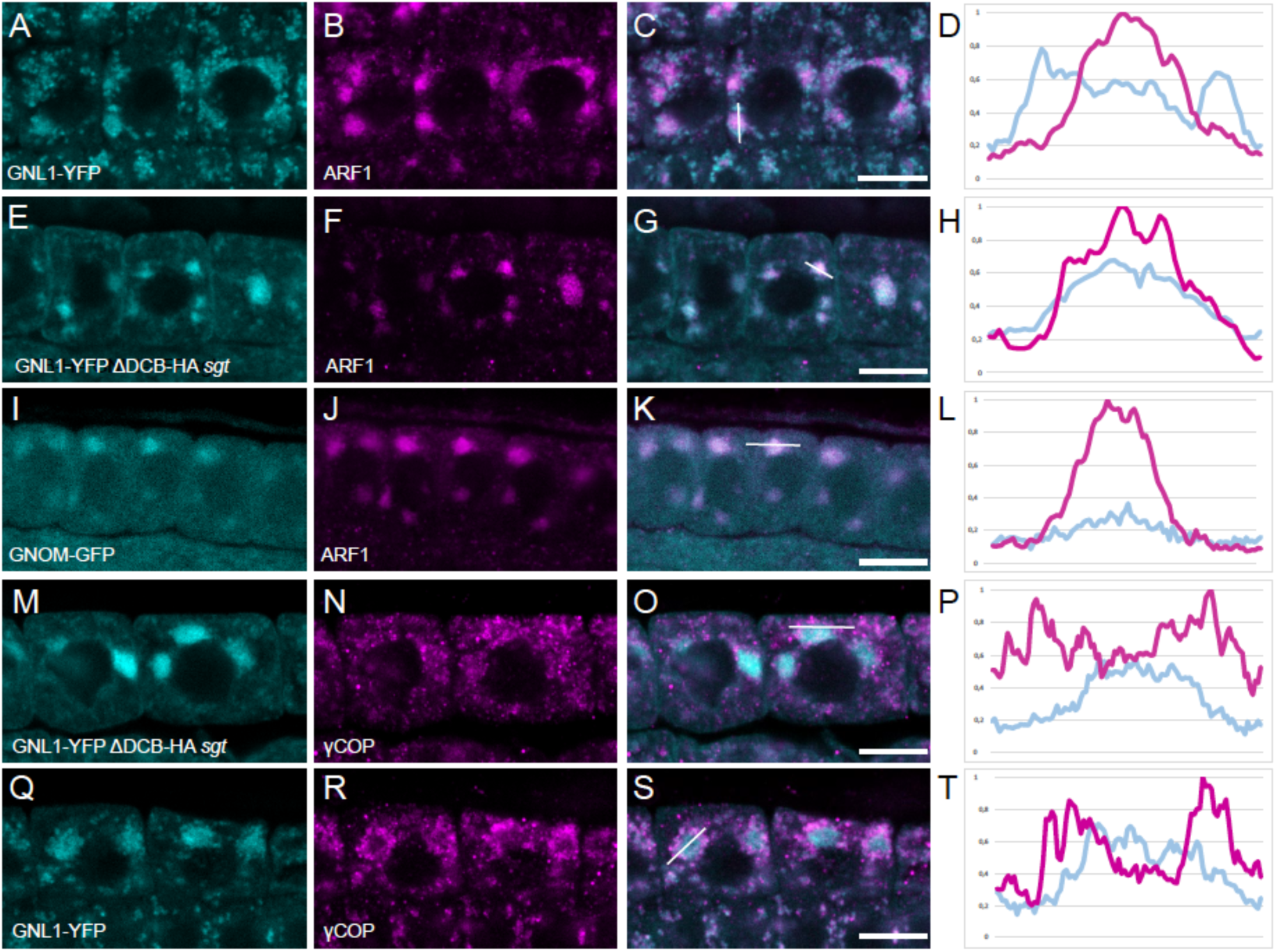
Subcellular relocation of Golgi-associated GNL1 to endosomes by interaction with ΔDCB^GNOM^. Seedlings expressing (**A**-**H**, **M**-**T**) GNL1-YFP or (**I**-**L**) GNOM-GFP were treated with 50 µM BFA for 1 h before fixation and immunostaining with anti-ARF1 antiserum (**A-L**, magenta) or anti-γCOP antiserum (**M-T**, magenta). Line scans (right panels) as indicated by the white lines in the adjacent panels. *GNOM* genotypes: (**A**-**D**, **I**-**L**, **Q**-**T**) wild-type; (**E**-**H**, **M**-**P**) *sgt* (*GNOM* and 4 adjacent genes on either side deleted) expressing ΔDCB^GNOM^. Note shift of GNL1 from γCOP-positive Golgi stacks to ARF1-positive BFA compartment caused by absence of DCB^GNOM^.

## Discussion

The paralogous ARF-GEFs GNOM and GNL1 are functionally divergent, with only GNOM required for polar recycling of auxin efflux carrier PIN1. Although they are expressed in the same cells, GNOM and GNL1 form homodimers but no heterodimers. To address the biological significance of preventing heterodimer formation, we engineered GNOM-GNL1 heterodimers, for example by deleting the N-terminal dimerisation domain of GNOM. The ΔDCB^GNOM^ fragment interacted with full-length GNL1, presumably via DCB-ΔDCB interaction required for membrane association as evidenced in the yeast two-hybrid interaction assay, and targeted the heterodimer to endosomes where GNOM normally acts. While the ΔDCB^GNOM^-GNL1 heterodimer was able to suppress the lethality of *gnom^sgt^* deletion mutant, lateral root development of the rescued seedlings was completely blocked. This deleterious effect demonstrated the necessity of keeping GNOM and GNL1 separate. In contrast to the ΔDCB^GNOM^-GNL1 heterodimer, the heterodimer consisting of chimeric DCB^GNL1^:ΔDCB^GNOM^ and full-length GNL1 had no such deleterious effect. However, the chimeric DCB^GNL1^:ΔDCB^GNOM^ protein was active on its own rather than dependent on its interaction with GNL1, as indicated by its rescue of the *gnom^sgt^ gnl1* double mutant. Thus, lateral root development of rescued *gnom^sgt^* seedlings was promoted by the separate activities of the chimeric DCB^GNL1^:ΔDCB^GNOM^ homodimers mediating endosomal recycling and GNL1 homodimers involved in COPI traffic required for secretion, although heterodimers consisting of chimeric DCB^GNL1^:ΔDCB^GNOM^ and full-length GNL1 also occurred. In conclusion, our observations suggest that coupling of GNOM-dependent recycling and GNL1-dependent secretion might be disadvantageous in competitive non-laboratory conditions, which would explain why the formation of GNOM-GNL1 heterodimers is normally prevented.

How is the formation of GNOM-GNL1 heterodimers prevented? Our results indicate that both GNOM and GNL1 form homodimers constitutively, i.e. only dimers but no monomers were detected both in the cytosol and on membranes. Thus, there seems to be a propensity of GNOM and GNL1 monomers to form (homo)dimers. The simplest assumption would be that the homodimers form during or immediately after protein synthesis, although direct evidence is lacking. Co-translational assembly of protein complexes has been reported before (Wells et al., 2015; Natan et al., 2017, 2018). In the case of GNOM, this precocious dimer formation is conceivable since the DCB domain of one GNOM protein interacts with the DCB domain of another GNOM protein and the DCB domain is located at the very N-terminus. Both GNOM and GNL1 use rare codons such that the rate of translation might be slow enough for folding of the DCB domain to occur while their translation is still ongoing. This “head start” of paired-up DCB^GNOM^ domains would facilitate their interactions with the two physically linked ΔDCB^GNOM^ fragments (Fig. 5A). In essence, this early homodimer formation would deplete the cell of GNOM monomers such that GNL1 monomers would be left to interact with one another to form (homo)dimers and thus, the formation of deleterious GNOM-GNL1 heterodimers might be prevented.

**Figure 5.**
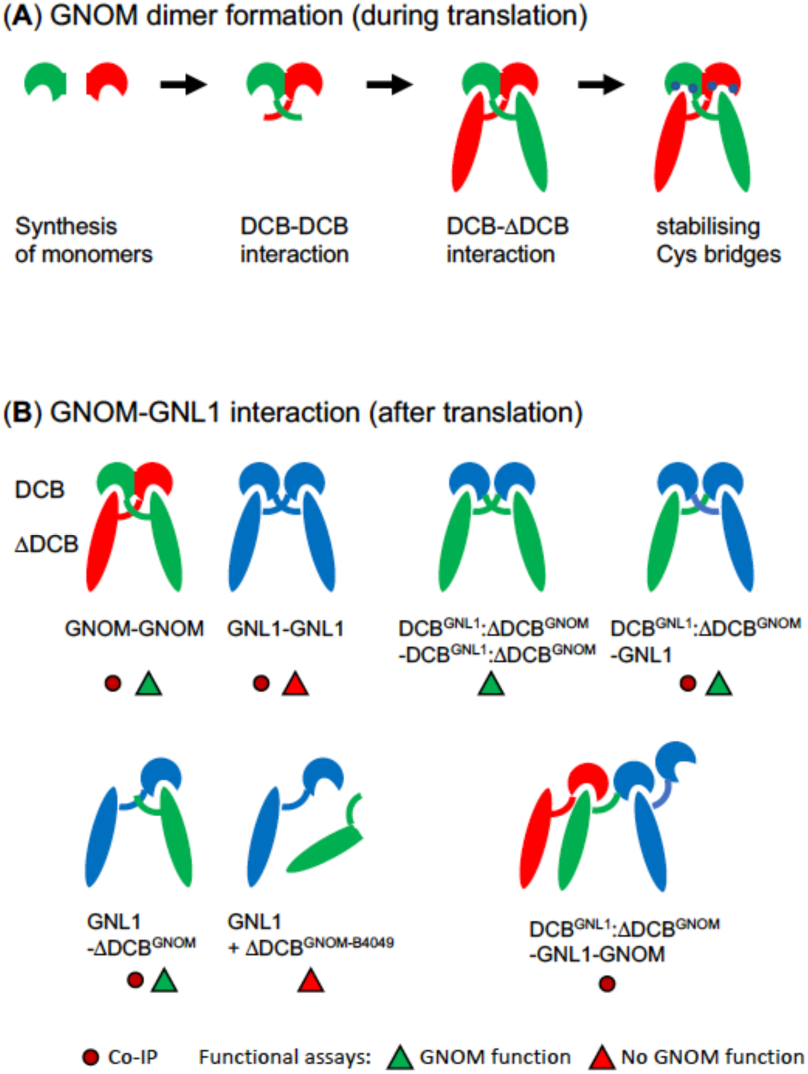
Role of DCB domain in GNOM dimer formation and GNOM-GNL1 interaction (model) (**A**) Stepwise GNOM dimer formation during translation. Interaction between two N-terminal DCB domains initiates dimer formation. The pair of fully translated proteins undergoes two DCB-ΔDCB interactions followed by the formation of stabilising Cys bridges (blue dots). (**B**) GNOM-GNL1 interactions established after translation. GNL1 and a chimeric DCB^GNL1^:ΔDCB^GNOM^ protein with GNOM function form dimers only mediated by DCB-ΔDCB interactions. Chimeric DCB^GNL1^:ΔDCB^GNOM^ protein can also act as a bridge between GNOM and GNL1 that do not interact directly. GNL1 interacts with ΔDCB^GNOM^ but not the mutant variant ΔDCB^GNOM-B4049^ that cannot interact with the DCB domain.

Our model predicts that if no initial DCB-DCB interaction takes place as in DCB^GNL1^:ΔDCB^GNOM^ chimeric protein or ΔDCB^GNOM^ fragment, there will be opportunity for interaction of the GNOM variant with GNL1, resulting in the formation of heterodimers to some extent or even heterotrimers as in the case of GNOM-DCB^GNL1^:ΔDCB^GNOM^-GNL1 detected by co-IP (Fig. 5B; see Fig. 3C). The opportunity for heterodimer formation also explains the complementation of mutant or truncated GNOM variants in the presence of GNL1 (Fig. 5B). Furthermore, the specific features of the individual GNOM variants determine whether their interaction with GNL1 leads to restoration of GNOM function (Fig. 5B).

Once the GNOM homodimer is fully formed including the DCB-ΔDCB interactions, cysteine-cysteine bridges appear to stabilise DCB-DCB and DCB-ΔDCB interactions between the identical subunits (Fig. 5A). This stabilisation might preclude the subsequent exchange of subunits between different GNOM homodimers as well as with GNL1 homodimers. Although Cys-to-Ser substitutions introduced into the DCB domain of GNOM appeared to affect DCB-DCB interactions more strongly than DCB-ΔDCB interactions, they seems to have general destabilising effects. It is thus likely that the Cys-to-Ser substitutions might only affect the stability, but not the initial formation, of the DCB-DCB interaction and thus do not interfere with precocious GNOM homodimer formation.

The mechanism proposed here for preventing GNOM-GNL1 heterodimer formation contrasts with the situation in bacteria where histidine kinases only form homomers to prevent cross-signalling, and this is mediated by a dimerisation domain that in each paralogue interacts with itself but not with its counterpart in other paralogues (Ashenberg et al., 2011).

Nonetheless, preventing heterodimers of functionally divergent paralogues has comparable effects in the two systems: like bacterial signalling pathways, plant trafficking pathways such as secretion and recycling can be regulated independently to meet specific challenges.

## Materials and Methods

### Plant genotypes and growth conditions

Columbia-0 (Col-0) and Landsberg *erecta* (Ler) were the *Arabidopsis thaliana* wild-type accessions used. The following mutant genotypes have been described previously: *gnom* alleles *emb30* and *B4049* (Busch et al., 1996), *gnom^sgt^* deletion (Brumm et al., 2020), *gnl1* T-DNA insertion (Richter et al., 2007), transgenic lines *GNOM-Myc*, *GNOM^ML^-Myc* and *GNOM-GFP* (Geldner et al., 2003), *ΔDCB^GNOM-HA^*, *ΔDCB^GNOM-Myc^* and *XLIM-ΔDCB^GNOM-B4049^-Myc* (Anders et al., 2008), *GNL1-YFP*, *GNL1-Myc* and *GNL1^LM^-Myc* (Richter et al., 2007). *ΔDCB^GNOM-HA^*, *ΔDCB^GNOM-Myc^* and *XLIM-ΔDCB^GNOM-B4049^-Myc* were again transformed into Col-0 and then crossed into the *gnom^sgt^* background in order to generate more independent transgenic lines.

Plants were grown on soil or agar plates under permanent light conditions (Osram L18W/840 cool white lamps) at 23°C and 40% humidity in growth chambers.

### Binary vector constructs, generation of transgenic plants, PCR genotyping and crosses

XLIM-*Δ*DCB^GNOM-B4049^-Myc, *Δ*DCB^GNOM^-HA, *Δ*DCB^GNOM^-Myc were crossed and/or transformed into heterozygous *gnom^sgt^/GNOM* and *gnom^sgt^/GNOM gnl1/GNL1* double mutant and analysed for complementation. Of three independent transgenic lines with good expression, one was chosen for further analysis. For co-immunoprecipitation analysis and whole-mount immunofluorescence staining, GNL1-Myc or GNL1-YFP were crossed with *Δ*DCB^GNOM^-HA in the *gnom^sgt^* mutant background.

To generate the DCB^GNL1^:ΔDCB^GNOM^ chimera, the DCB domain of GNL1 was amplified via primer extension PCR and inserted, via PmeI und SwaI restriction sites, into the genomic fragment GNXbaI^wt^-myc (Geldner et al., 2003) in pBlueScript. The following primers were used:

GN_DCB_UP_F 5’ TCGTTCTAGCGTCGAACAAACTCCTCGTTTTCTTTGATTCGCATTG 3’
GN_DCB_UP_R 5’ CCCGAAGGATGATTCTGATACCCCATTTAATCTGCTCAAATCTTCA 3’
GNL1_DCB_S 5’ TGAAGATTTGAGCAGATTAAATGGGGTATCAGAATCATCCTTCGGG 3’
GNL1_DCB_AS 5’ TCTGTTCTTTCAACATCTGGAAGTTGAGAGAAGATACATCTAATC 3’
GN_DCB_DW_F 5’ GATTAGATGTATCTTCTCTCAACTTCCAGATGTTGAAAGAACAGA 3’
GN_DCB_DW_R 5’ TGGTCTAAATTCTCTAGTAGTGATCTTTGACTCGCACTAGGAAAA 3’

The pGN:DCB^GNL1^:*Δ*DCB^GNOM^-Myc fragment was first inserted into an intermediate pBar vector via XbaI restriction sites and afterwards introduced into pGII(BAR) expression vector and transformed into Col-0 background. T1 plants were selected using phosphinotricine.

Lines showing good expression were crossed with heterozygous *gnom^sgt^/GNOM*, *gnl1/GNL1* and *gnom^sgt^/GNOM gnl1/GNL1* double mutants. For co-immunoprecipitation analysis, transgenic plants expressing DCB^GNL1^:*Δ*DCB^GNOM^-Myc from *GNOM* regulatory sequences were crossed with plants bearing the transgenes *GNOM-GFP* or *GNL1-YFP*.

To generate *GNOM* coding sequences with the desired C-to-S mutations (GNOM^xCS^) for plant expression, the respective DCB^GNOM-xCS^ fragment was amplified from the yeast vector (*pJG4-5-DCB^GNOM-xCS^*) using the following primers:

GNDCB_YV_F: 5’ CTCCCGAATTCGCAGATTTAATGGGTCGCCTA 3’
GNDCB_YV_R. 5’GCTTCTCGAGCTATTGTTTGATGCTAC 3’.

The purified PCR product was used as primer pair for site-directed mutagenesis of *pDONOR221-GNOM* to obtain *pDONOR221-GNOM^xCS^*. The yeast vector harbouring *DCB^GNOM^* and *DCB^GNOM-xCS^* had an un-annotated V210I mutation, which was repaired by site-directed mutagenesis of *pDONOR221-GNOM^xCS^* plasmid using the following primers:

GNOM_DCB_Ile-Val_F: 5’ GTTATTGCAACGAGTAGCTCGCCACACGATGCA
GNOM_DCB_Ile-Val_R: 5’ CGTGTGGCGAGCTACTCGTTGCAATAACTCA.

The sequence corresponding to the N-terminal part of *GNOM^xCS^* (1-682 amino acids) was amplified from *pDONOR221-GNOM^xCS^* plasmid and cloned into a pre-existing *pGII(Bar)-pGNOM:GNOM-YFP* plasmid using PspXI (New England Biolabs catalogue no R0656) and MscI (Thermo Scientific catalogue no ER1211) enzymes to generate *pGII(Bar)-pGNOM:GNOM^xCS^-YFP*. The following primers were used for amplification of *GNOM^xCS^* (1-682 amino acids):

DCB_CS_PspXI_LinkerAvrII: 5’ CTTCCTCGAGGTCCTAGGACATGGGTCGCCTAAAGTT 3’
pGIIGNOM_DCB_CS_MscI: 5’ CCTTTGGCCAGAATCTCAGGAGATTGCATATAGTA 3’

To generate *pGII(Bar)-pGNOM:GNOM^4CS^-Myc*, the *YFP* sequence in *pGII(Bar)-pGNOM:GNOM^4CS^-YFP* was replaced by a *3xMyc* sequence using SmaI and XbaI restriction sites.

The binary vectors were transformed into Arabidopsis wild-type (Col-0) and T1 plants were selected using BASTA (Bayer catalogue no 79011725). T1 Plants showing good expression were used for crossing with *gnom^sgt^/GNOM*, *gnom^sgt^/GNOM gnl1/GNL* or other transgenic lines.

Genotyping of *gnom^sgt^* was performed using the following primers:

GN_overtag_S: 5’ GAAAGTGAAAGTAAGAGGC 3’
GN_overtag_AS: 5’ CGTAGAGAGGTGTTACATAAG 3’

Genotyping of *gnl1* was performed as described earlier (Richter et al., 2007).

### Yeast two-hybrid interaction assays

DCB^GNOM^ (aa1-246), ΔDCB^GNOM^ (aa232-1451), ΔDCB^GNOM-B4049^ (aa232-1451; G_579_R, *B4049* mutation) and ΔDCB^GNOM-HUS-BOX^ (aa232-1451; D_468_G, mutation in HUS box) constructs and assay were as described (Grebe et al., 2000; Anders et al., 2008).

DCB^GNL1^ (aa1-244) was cloned into standard yeast two-hybrid vectors pEG202 and pJG4-5 via PCR-introduced EcoRI and XhoI restriction sites. ΔDCB^GNL1^ (aa245-1443) was cloned into modified pEG202 and pJG4-5 vectors (in which the EcoRI restriction site in the MCS was replaced by a NotI site; designated pM8 and pM5; Grebe et al., 2000) via NotI and XhoI. DCB^GNOM-1CS^ was synthesized by the company BaseGene B.V. (Leiden, Netherlands) and then cloned into pEG202 and pJG4-5 via EcoRI and XhoI.

The generation of DCB^GNOM:GNL1^ chimeras was based on subdivision of the DCB domains into four fragments, each representing roughly one quarter of the domain (Figure 1 – figure supplement 1): For DCB^GNOM^ the first quarter comprises base pairs (bp) 1-183/ amino acids (aa) 1-61; the second quarter bp 184-432 / aa 62-144; the third quarter bp 433-528 / aa 145-176,; and the fourth quarter bp 529-672 / aa 177-224. For DCB^GNL1^ the subdivision is as follows: bp 1-177 / aa 1-59; bp 178-426 / aa 60-142; bp 427-528 / aa 143-176; bp 529-732 / aa 177-244. The respective borders for these “quarters” were chosen based on a sequence alignment between DCB^GNOM^ and DCB^GNL1^.

Chimeras 1, 2, 3, and 4 were generated via PCR. In two individual PCR reactions on GNOM and GNL1 templates, an N-terminal and a C-terminal fragment comprising the respective number of quarters were created. Overlaps between them were introduced on one side of each fragment through primer extension. A third PCR using these overlapping fragments as templates produced the unified chimeric DCB sequence.

Chimera 5 was generated using chimera 4 and GNL1 as templates for the first two PCRs, and chimera 6 was generated using chimera 1 and GNOM as templates, followed by the joining PCRs.

All chimeric DCB sequences were then cloned into pEG202 and pJG4-5 via EcoRI and XhoI restriction sites. The restriction enzymes were supplied by New England Biolabs and have the following catalogue numbers: EcoRI (R3101S), XhoI (R0146L), NotI (R3189L).

The following primers were used:

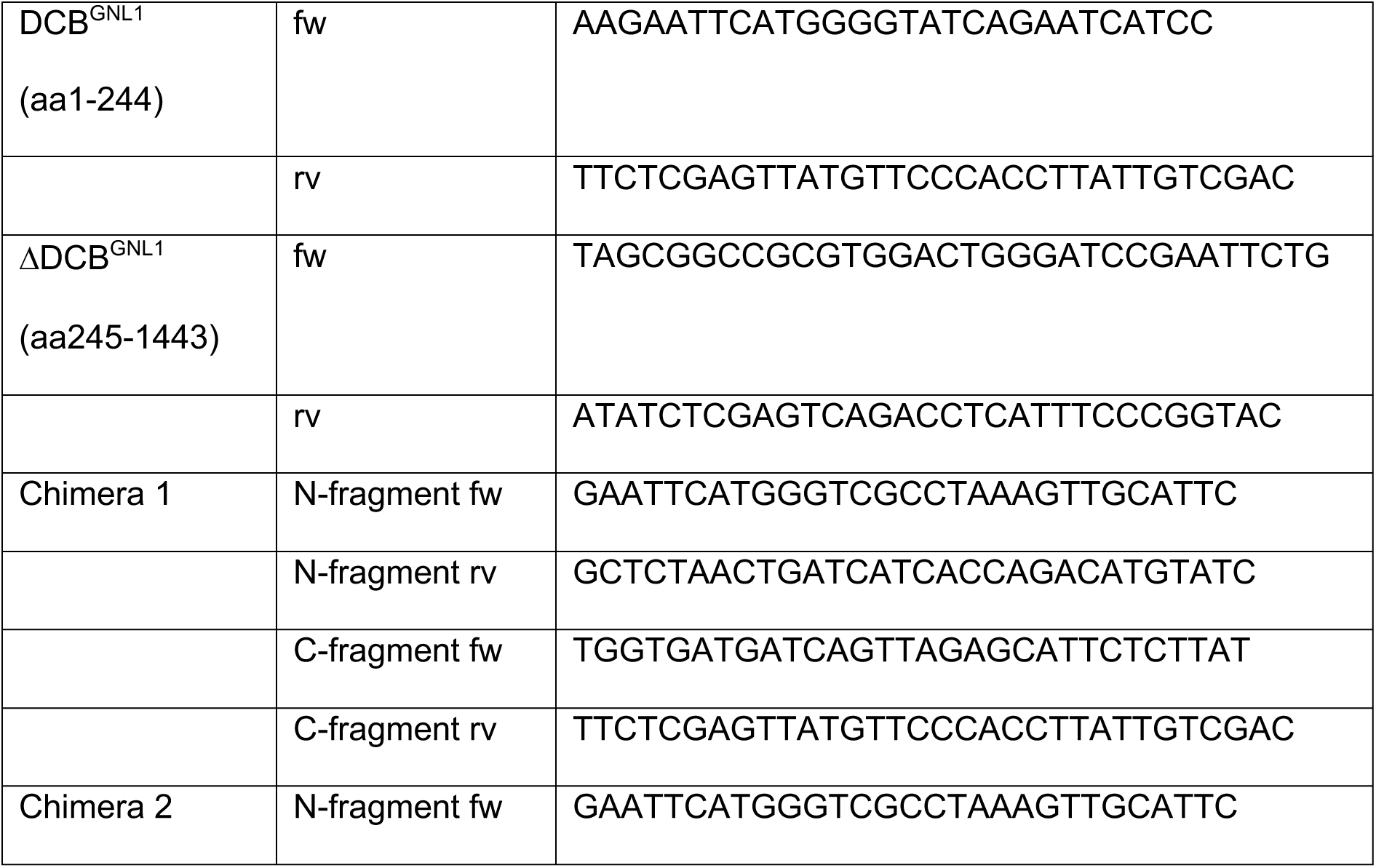
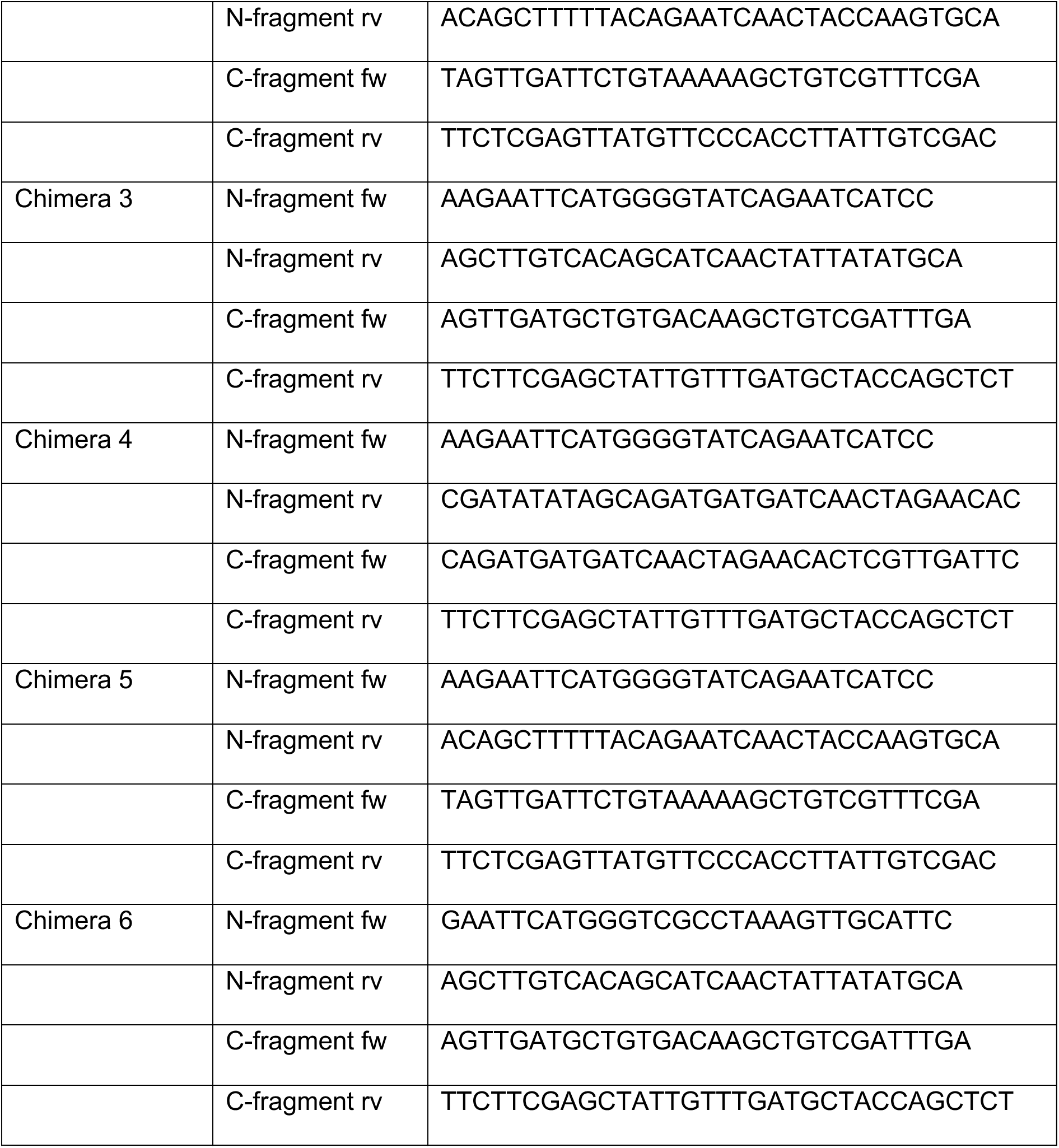

To introduce C-to-S mutations into DCB^GNOM^, primer-based mutagenesis was performed on *pJG4-5-DCB^GNOM^* plasmid as template. PCR products carrying different C-to-S mutations were combined by primer extension PCR to generate *DCB^GNOM-7CS^*, *DCB^GNOM-4CS^*, and *DCB^GNOM-3CS^* and cloned in *pEG202* and *pJG4-5* vectors using EcoRI and XhoI. The following primers were used for DCB^GNOM^ mutagenesis and cloning:

GNDCB_YV_F: 5’ CTCCCGAATTCGCAGATTTAATGGGTCGCCTA 3’
GNDCB_C22S_R: 5’ AGTGGTTGTATTACTTGAATCAGTACTCTCAAAG 3’
GNDCB_C33S_F: 5’ GATTCAAGTAATACAACCACTTTAGCAAGCATGA 3’
GNDCB_C148S_R: 5’ CTCAAATCGACTGCTTGTCACAGA 3’
GNDCB_C148S_F: 5’ TCTGTGACAAGCAGTCGATTTGAGGTG 3’
GNDCB_C172S_R: 5’ GATGCTTTATTTTTCATACTTGCTAGAAGAAC 3’
GNDCB_C172S_F: 5’ GTTCTTCTAGCAAGTATGAAAAATAAAGCATC 3’
GNDCB_C187&193S_R: 5’ GAAAACTAGTGTTGACGACAGTGCTTACATG 3’
GNDCB_C187&193S_F: 5’ CATGTAAGCACTGTCGTCAACACTAGTTTTC 3’
GNDCB_C221S_R: 5’ GATGCGAGAAGATACTCCTCACTAATTC 3’
GNDCB_C221S_F: 5’ GAATTAGTGAGGAGTATCTTCTCGCATC 3’
GNDCB_S172C_R: 5’ GATGCTTTATTTTTCATACATGCTAGAAGAAC 3’
GNDCB_S172C_F: 5’ GTTCTTCTAGCATGTATGAAAAATAAAGCATC 3’
GNDCB_YV_R: 5’ GCTTCTCGAGCTATTGTTTGATGCTAC 3’

### Yeast two-hybrid – quantitative oNPG assays

The quantitative oNPG assays shown in Figure 1D-H and Figure 1 – figure supplement 1 were done as follows. For each tested combination, six biological replicates (yeast cultures) were harvested after determining OD600, and then each subdivided into two technical replicates. After measurement of absorption at 420nm, mean values for each pair of technical replicates were calculated and then used to calculate the ß-galactosidase activity of the biological replicates, of which again a mean value as well as standard deviation was calculated. Of the six determined ß-galactosidase activity values for each interactor combination, the highest and lowest values were excluded from the calculation of the mean activity, resulting in a sample size of n=4. The experiment shown in Fig. 1E was performed three times with similar results, the other ONPG assays once. The yeast cultures were randomly picked and inoculated. The scientists performing the experiments were aware of sample identity.

### Physiological assays

Root gravitropic response of 50 five-days old seedlings was measured by ImageJ software after transferring seedlings to BM plates (no brefeldin A, BFA) or agar plates containing 10 μM BFA (Sigma catalogue no: B7651) and rotating the plates vertically by 135° for 24h (Richter et al., 2007) . Lateral root primordia formation was analysed after transferring 7-days old seedlings to 5μM 1-naphthaleneacetic acid (NAA)-containing liquid MS medium and clearing the roots after treatment overnight or for 2 days (Geldner et al., 2004). Light microscopy images were taken with a Zeiss Axiophot microscope, Axiocam and AxioVision_4 Software. Image size, brightness and contrast were edited with Adobe Photoshop CS 3 Software.

For each assay, the experiment was repeated at least twice and the numbers of seedlings analysed are indicated as n in Fig. 3B, D and E. Fig. 3D shows data from one of 4 experiments, Fig. 3E from one of 6 experiments. Figure 3 – figure supplement 2 shows images of at least 5 different seedlings from 1 of 3 experiments. In each experiment, 10-20 seedlings were analysed.

Post-embryonic phenotypes shown in Figure 2 – figure supplements 2 and 5, and Figure3 – figure supplement 1 were from 2 or 3 experiments each involving at least 5 different plants of the relevant genotype.

### Live-cell imaging and whole-mount immunofluorescence staining

Live-cell imaging of 5-days old Arabidopsis seedlings was performed after 1h treatment with 50 μM BFA (Sigma catalogue no: B7651) and 2 μM FM4-64 (SynaptoRed C2, Sigma catalogue no S6689).

For whole-mount immunofluorescence staining, four to six-days old seedlings were incubated in 24-well cell-culture plates in 50 μM BFA-containing liquid growth medium (0.5x MS medium, 1% sucrose, pH 5.8) (Duchefa Biochemie catalogue no M0221.005) at 23°C for 1 hour and then fixed in 4% paraformaldehyde in MTSB at room temperature for 1 hour.

Whole-mount immunofluorescence staining was performed manually (Lauber et al., 1997) or with an InsituPro machine (Intavis; Müller et al., 1998). All antibodies were diluted in 1x PBS buffer. The following antisera were used for immunofluorescence staining: rabbit polyclonal anti-ARF1 (Agrisera AS08 325) diluted 1:1000; rabbit polyclonal anti-AtγCOP (Agrisera AS08 327) diluted 1:1000; goat anti-rabbit CY3-conjugated secondary antibodies (Dianova catalogue no 111-165-144) were diluted 1:600. Nuclei were stained with 4′,6-diamidino-2-phenylindole (DAPI; Sigma-Aldrich catalogue no D9542, 1:600 dilution). Figure 4 shows images of seedlings from 1 of 6 experiments. In each experiment, 20-40 seedlings were mounted for immunostaining and at least 10 roots were analysed per genotype.

### Confocal microscopy and processing of images

Fluorescence images were acquired at the confocal laser scanning microscope TCS-SP8 from Leica or LSM880 from Zeiss, using a 63x water-immersion objective and Leica or Zeiss software (Leica LAS X; Zeiss Zen), respectively. Overlays and contrast/brightness adjustments of images were performed with Adobe Photoshop CS3 software. Intensity line profiling was performed with Leica software (LAS X).

### Co-immunoprecipitation analysis

The immunoprecipitation protocol was modified from Singh et al. (2014). Specifically, 0.5-3g of 8 to 10-days old Arabidopsis seedlings were homogenized in 1:1 lysis buffer (50mM Tris pH 7.5, 150mM NaCl, 2mM EDTA) containing 1% Triton-X100 and protease inhibitors (cOmplete EDTA-free®, Roche catalogue no 04693132001). For immunoprecipitation, anti-Myc-agarose beads (Sigma catalogue no A7470) or anti-HA-agarose beads (Sigma catalogue no A2095) or GFP-Trap beads (Chromotek catalogue no gta20) were incubated with plant extracts at 4°C for 2h30min. Beads were then washed twice with wash buffer containing 0.1% Triton-X100 and 1-2 times without Triton-X100. Bound proteins were eluted by boiling the beads in 2x Laemmli buffer at 95°C for 5min. All co-immunoprecipitation experiments were repeated at least twice, except for the trimer co-IP shown in Fig. 3C.

### Subcellular fractionation

Subcellular fractionation was performed as described (Brumm et al., 2020). Briefly, 3-4 g of Arabidopsis seedlings were ground in liquid nitrogen, suspended in 2x volume of lysis buffer (50mM Tris pH 7.5, 150mM NaCl, 2mM EDTA) supplemented with protease inhibitors (cOmplete EDTA-free®, Roche catalogue no 04693132001) and centrifuged at 10,000 x *g* for 15 min at 4°C. The supernatant (S10) was subjected to 100.000 x *g* centrifugation for 1h at 4°C. The pellet (P100) was suspended in extraction buffer containing 1% Triton-X100 (v/v) and solubilized by sonication. The supernatant (S100) was also supplemented with Triton-X100, to a final concentration of 1%. The subcellular fractionation experiment was not repeated as it was also performed in Brumm et al. (2020) and no unexpected result was obtained here.

### Sample preparation for mobility shift assay in SDS-PAGE

Soluble protein extracts were prepared similar to immunoprecipitation experiments, frozen in liquid N2 and stored at -80^°^C until further use. At the time of loading on SDS-PAGE, protein extracts were thawed on ice, mixed 1:1 with Laemmli buffer with or without reducing agent (5% ß-mercaptoethanol, BME; Carl Roth catalogue no 4227.3, or dithiothreitol, DTT; Carl Roth catalogue no 6908.1) and boiled at 95°C for 5min.

### SDS-PAGE and protein gel blotting

SDS-PAGE gel electrophoresis and protein gel blotting with PVDF membranes (Thermo Scientific catalogue no 88520) were performed as described (Lauber et al., 1997). All antibodies were diluted in 5% milk/TBS-T solution. Antibodies and dilutions: mouse anti-c-Myc mAB 9E10 (Santa Cruz Biotechnology catalogue no sc-40), 1:1000; mouse anti-GFP (Roche catalogue no 11814460001), 1:2500; mouse anti-LexA mAb C-11 (Santa Cruz Biotechnology sc-390386), 1:1000; POD-conjugated anti-HA (Roche catalogue no 2013819), 1:4000; rabbit anti-SEC7^GNOM^ (Steinmann et al., 1999), 1:2500; rabbit anti-SEC21 antiserum (Agrisera catalogue no AS08 327; Pimpl et al., 2000), 1:2000; anti-mouse (Sigma catalogue no A2554) or anti-rabbit peroxidase-conjugated (Merck Millipore catalogue no AP307P) or alkaline phosphatase-conjugated antibodies (Jackson Immuno Research catalogue no 111-055-003), 1:5000. Detection was performed with the BM-chemiluminescence blotting substrate (Roche catalogue no 11500708001) and FusionFx7 imaging system (PeqLab). Image assembly was performed with Adobe Photoshop CS3.

## Acknowledgements

We thank Marika Kientz for expert technical assistance and discussions, and Steffen Lau for the initial cloning of the DCB domain of GNL1 in the yeast B42 vector. This study was funded by the Deutsche Forschungsgemeinschaft (grant SFB1101/TPA01 to G.J.).

## Author Contributions

SB, MKS, HB, SR and GJ conceived the idea and designed the experiments. SB, MKS, HB, SR, KH, TK, SBa, CK, HW and ST performed the experiments. SB, MKS, TK, HW and ST cloned constructs for in-vivo analysis, generated transgenic lines and performed genetic analyses. SB performed microscopic imaging analyses, MKS analysed protein interaction by co-immunoprecipitation, and HB, MKS, TK and SBa performed yeast two-hybrid interaction studies. SB and GJ wrote the manuscript with input from all authors.

---

**Figure 1 – figure supplement 1.**
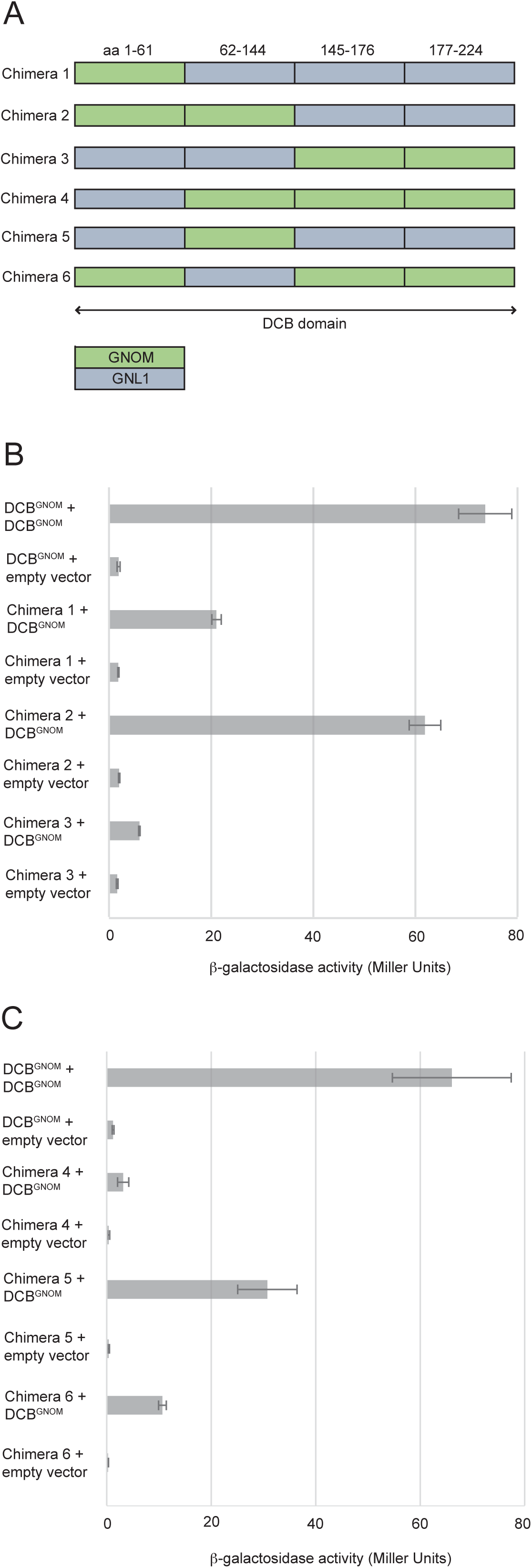
DCB-DCB interaction assays of GNOM-GNL1 chimeric DCB domains. Yeast two-hybrid assays revealed that the chimeric DCB domain #2 with aa1-144 from GNOM displayed nearly full interaction with DCB^GNOM^ whereas the reciprocal chimera #3 lost most of its interaction activity.

**Figure 2 – figure supplement 1.**
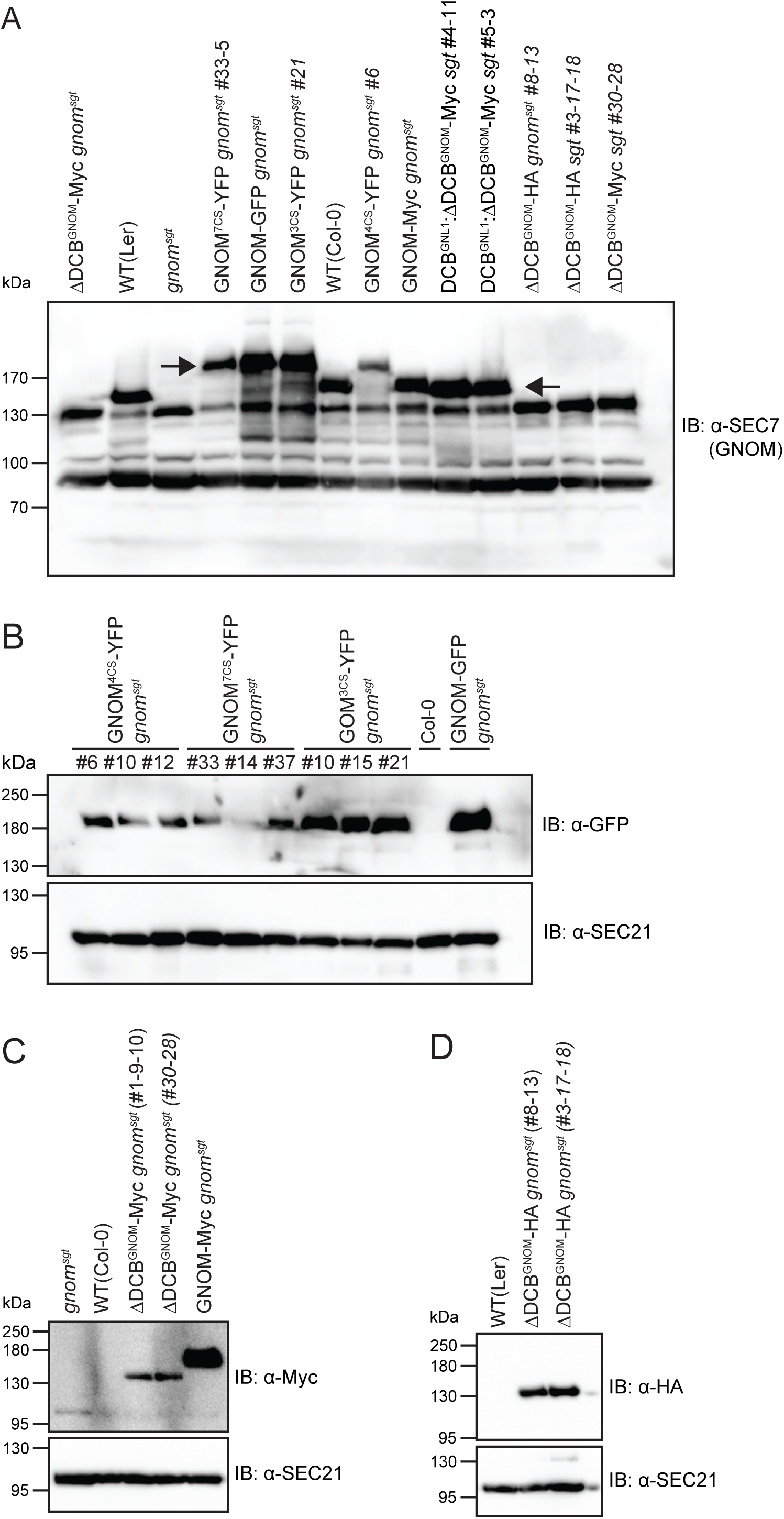
Expression of transgenes in *gnom^sgt^* background. (**A**) Protein extracts of *gnom^sgt^* mutant seedlings rescued by expression of different transgenes probed with anti-SEC7^GNOM^ antiserum. Arrows indicate GNOM bands. The cross-reacting band at approx. 90 kDa serves as an internal control. The band representing Myc-tagged or HA-tagged ΔDCB^GNOM^ at 130 kDa overlaps with a cross-reacting band present in all samples. (**B**) Protein extracts of *gnom^sgt^* mutant seedlings rescued by expression of YFP-tagged GNOM^CS^ substitution variants probed with anti-GFP antibody. Loading control: band detected with anti-SEC21 antiserum. (**C, D**) Protein extracts of *gnom^sgt^* mutant seedlings rescued by expression of (C) Myc-tagged or (D) HA-tagged ΔDCB^GNOM^. Controls: *gnom^sgt^,* GNOM deletion; WT(Col-0), wild-type; GNOM-Myc *gnom^sgt^*, Myc-tagged full-length GNOM expressed in *gnom^sgt^* deletion background. Loading control: band detected with anti-SEC21 antiserum.

**Figure 2 – figure supplement 2.**
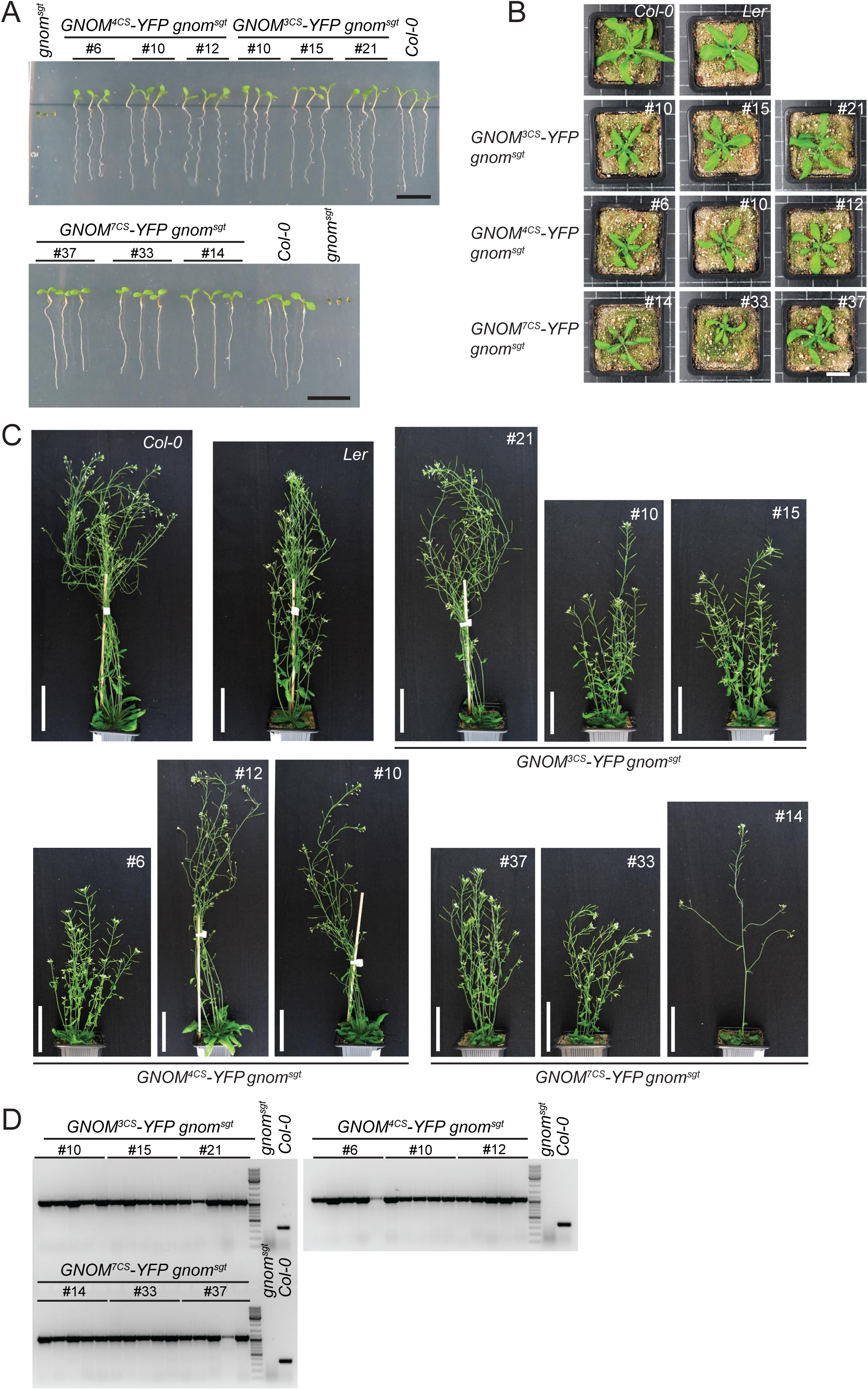
Rescue of *gnom^sgt^* mutant plants with C-to-S substitution variants of GNOM. (**A**) Seedling phenotypes. Scale bars, 1 cm. (**B, C**) Postembryonic phenotypes of *GNOM^CS^* transgenic plants: (B) rosette-stage plants on 23 dag; (C) adult plants on 38 dag. Scale bars, 2 cm (B), 6 cm (C). (**D**) PCR showing rescue of *gnom^sgt^* by different *GN^CS^-YFP* transgenes. Five seedlings from each line were genotyped using GN overtag primer.

**Figure 2 – figure supplement 3.**
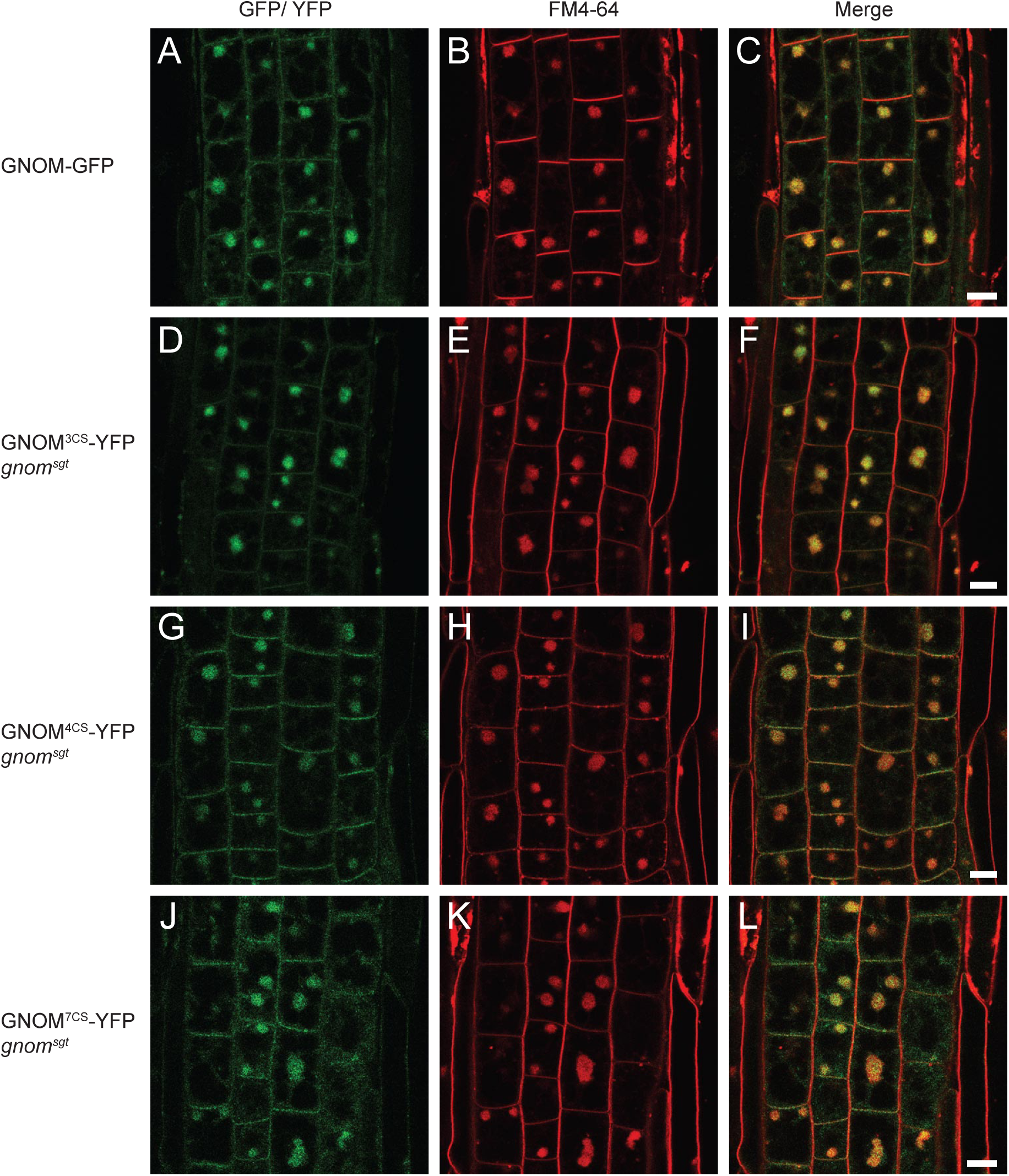
Subcellular localisation of GNOM^CS^ mutant proteins. YFP-tagged GNOM^CS^ proteins localised to BFA compartments stained with the endocytic tracer FM4-64 in root cells of seedlings treated with 50 µM BFA for 1h. GN-GFP (control) is in wild-type background and the YFP-tagged GNOM^CS^ variants are in the *gnom^sgt^* deletion mutant background. Scale bars, 10 μm.

**Figure 2 – figure supplement 4.**
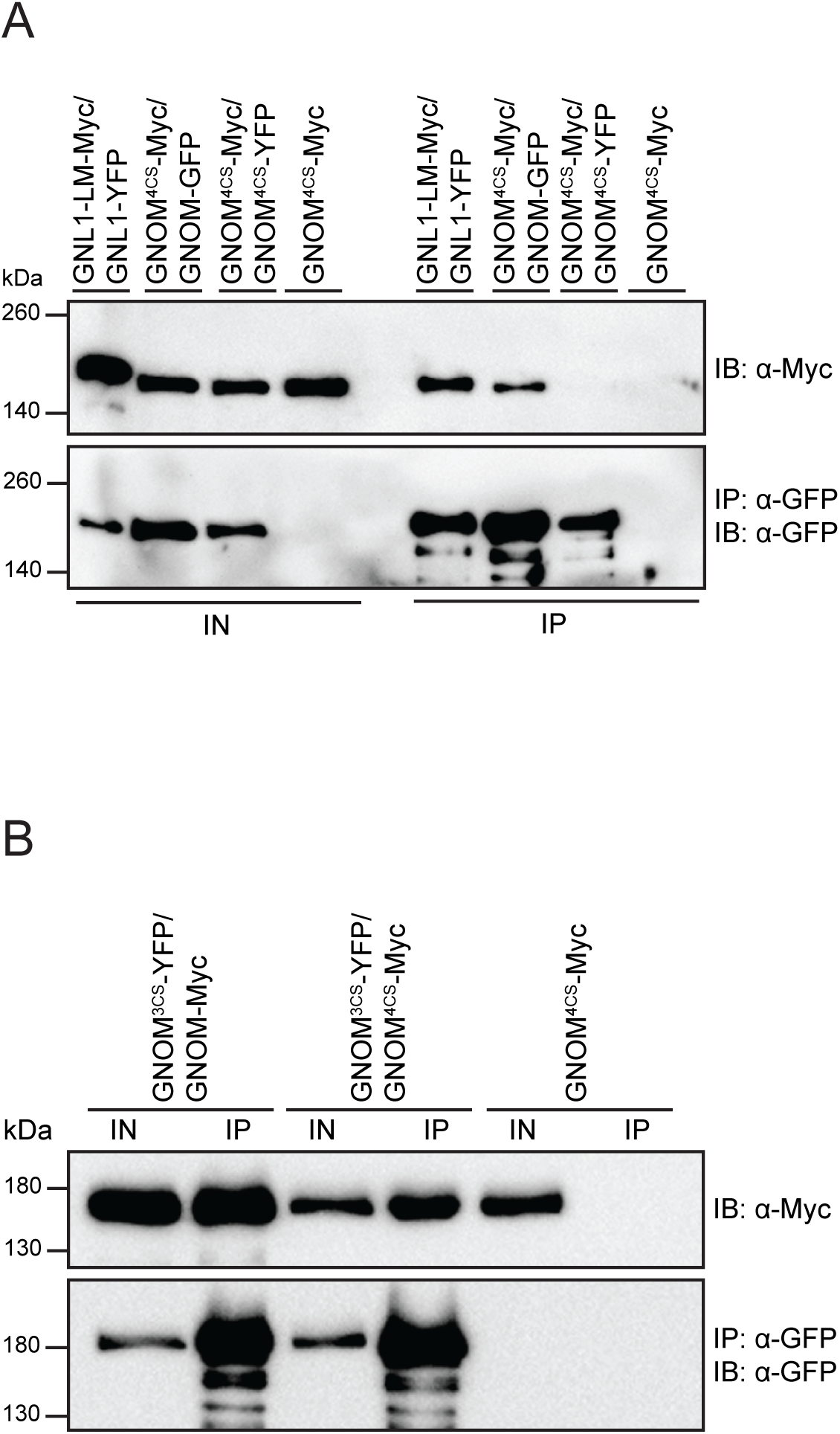
Interaction behaviour of GNOM^4CS^. Protein extracts of transgenic seedlings expressing differently tagged GNOM, GNL1, GNOM^4CS^ or GNOM^3CS^ proteins were subjected to co-immunoprecipitation analysis with anti-GFP beads. (**A**) GNOM^4CS^ interacted with GNOM, but failed to interact with itself (GNOM^4CS^-YFP). (**B**) GNOM^4CS^ interacted with GNOM^3CS^, which reflects the ability of DCB^GNOM-3CS^ to interact with ΔDCB^GNOM^, in contrast to the inability of DCB^GNOM-4CS^ (see Figure 2E). IB, immunoblot detection; IN, input; IP, immunoprecipitate. Size markers on the left (in kDa).

**Figure 2 – figure supplement 5.**
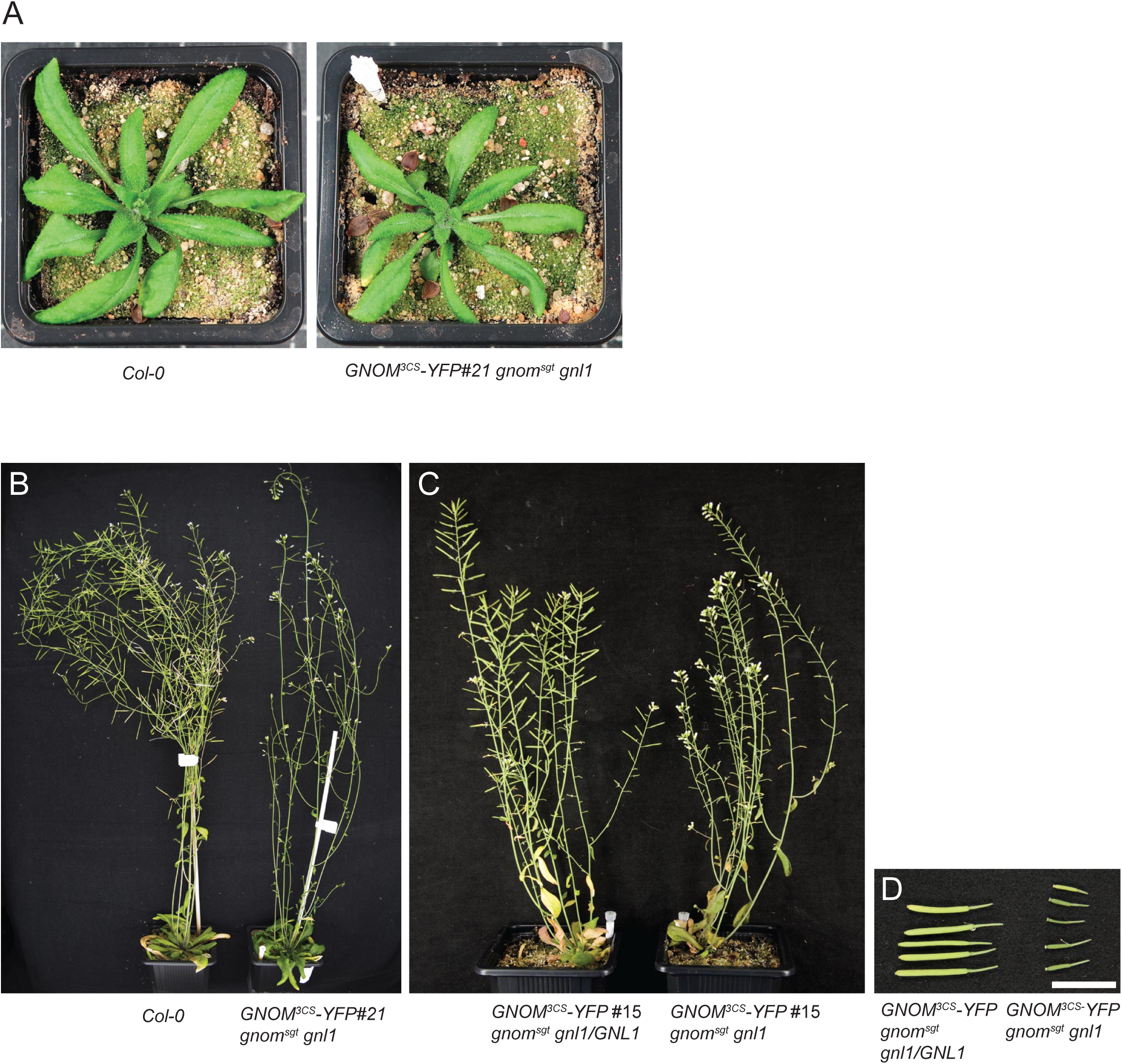
Complementation of *gnom^sgt^ gnl1* double knockout mutant with GNOM^3CS^. Although GNOM^3CS^ rescues development of the double knockout, giving rise to adult plants (**A-C**), the rescued double mutants are sterile as indicated by the small siliques without fertilised ovules (**D**). Col-0, wild-type control. Scale bar, 1 cm.

**Figure 3 – figure supplement 1.**
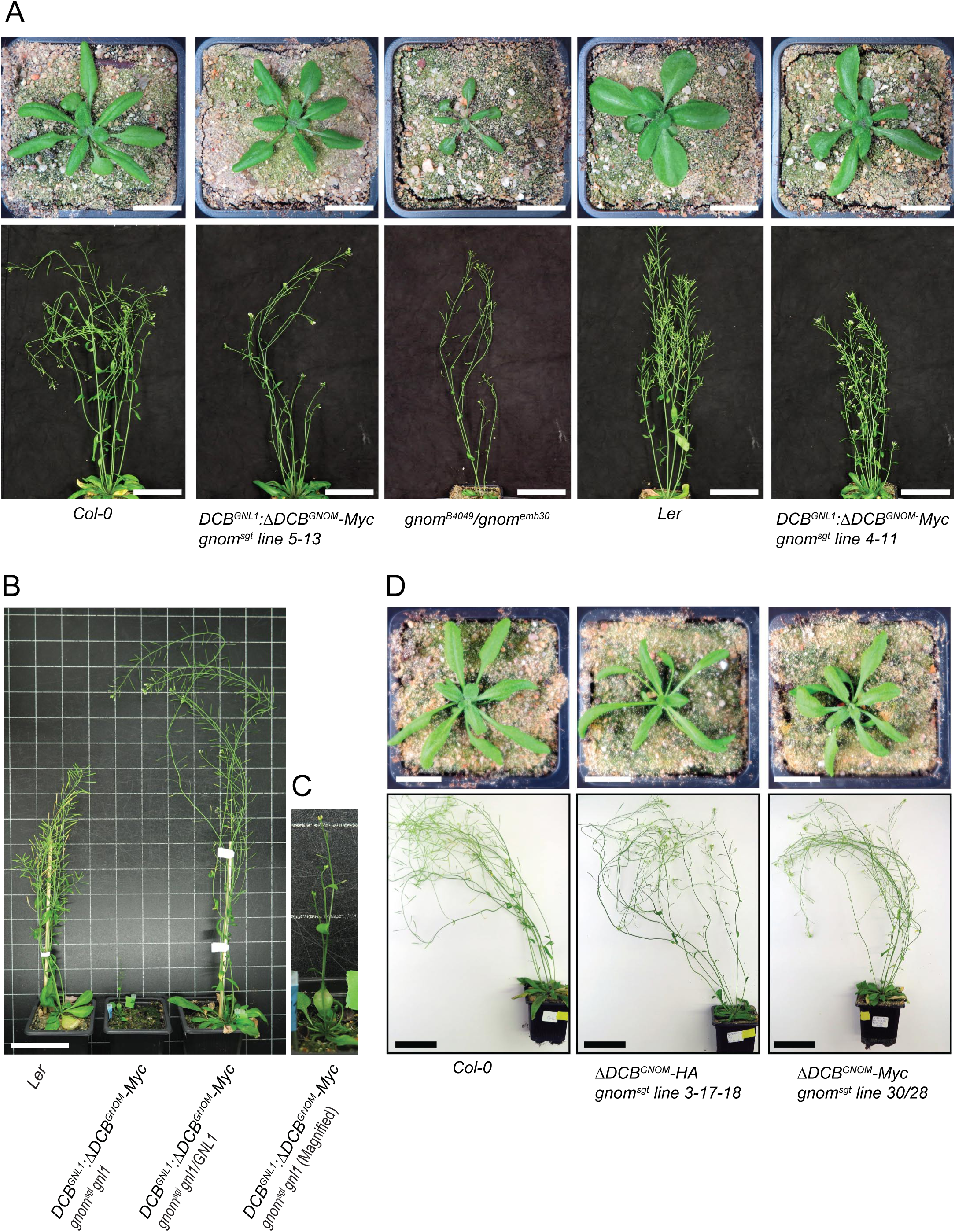
Postembryonic phenotypes of *gnom^sgt^* deletion mutant and *gnom^sgt^ gnl1* double mutant rescued by expression of chimeric DCB^GNL1^:ΔDCB^GNOM^ or ΔDCB^GNOM^ protein. (**A**) Two independent *DCB^GNL1^:ΔDCB^GNOM^* transgene insertions (#4-11 and #5-13) rescue the *gnom^sgt^* deletion mutant. *Top*: Three weeks old seedlings. Scale bars, 2 cm. *Bottom*: Six weeks old plants. Scale bars, 7 cm. Controls: Col-0, wild-type; Ler, wild-type (parental genotype of *gnom^sgt^* deletion mutant); *b4049/emb30,* complementing non-functional *gnom* alleles. (**B, C**) Rescue of *gnom^sgt^ gnl1* double mutant by expression of Myc-tagged DCB^GNL1^:ΔDCB^GNOM^ chimeric protein. Note strongly reduced stature of the rescued double mutant (**B**, middle; **C**, at higher magnification) as compared to the rescued *gnom^sgt^* single mutant (**B**, right). Ler, wild-type control. Scale bar, 7 cm. (**D**) Differently tagged *ΔDCB^GNOM^* transgenes are able to rescue the *gnom^sgt^* deletion mutant. *Top*: Three weeks old seedlings. Scale bars, 2cm. *Bottom*: Six weeks old plants. Scale bars, 7 cm. Col-0, wild-type; *ΔDCB^GNOM^-HA gnom^sgt^* and *ΔDCB^GNOM^-Myc gnom^sgt^*, *gnom* deletion mutant expressing HA-tagged or Myc-tagged ΔDCB^GNOM^ fragment.

**Figure 3 – figure supplement 2.**
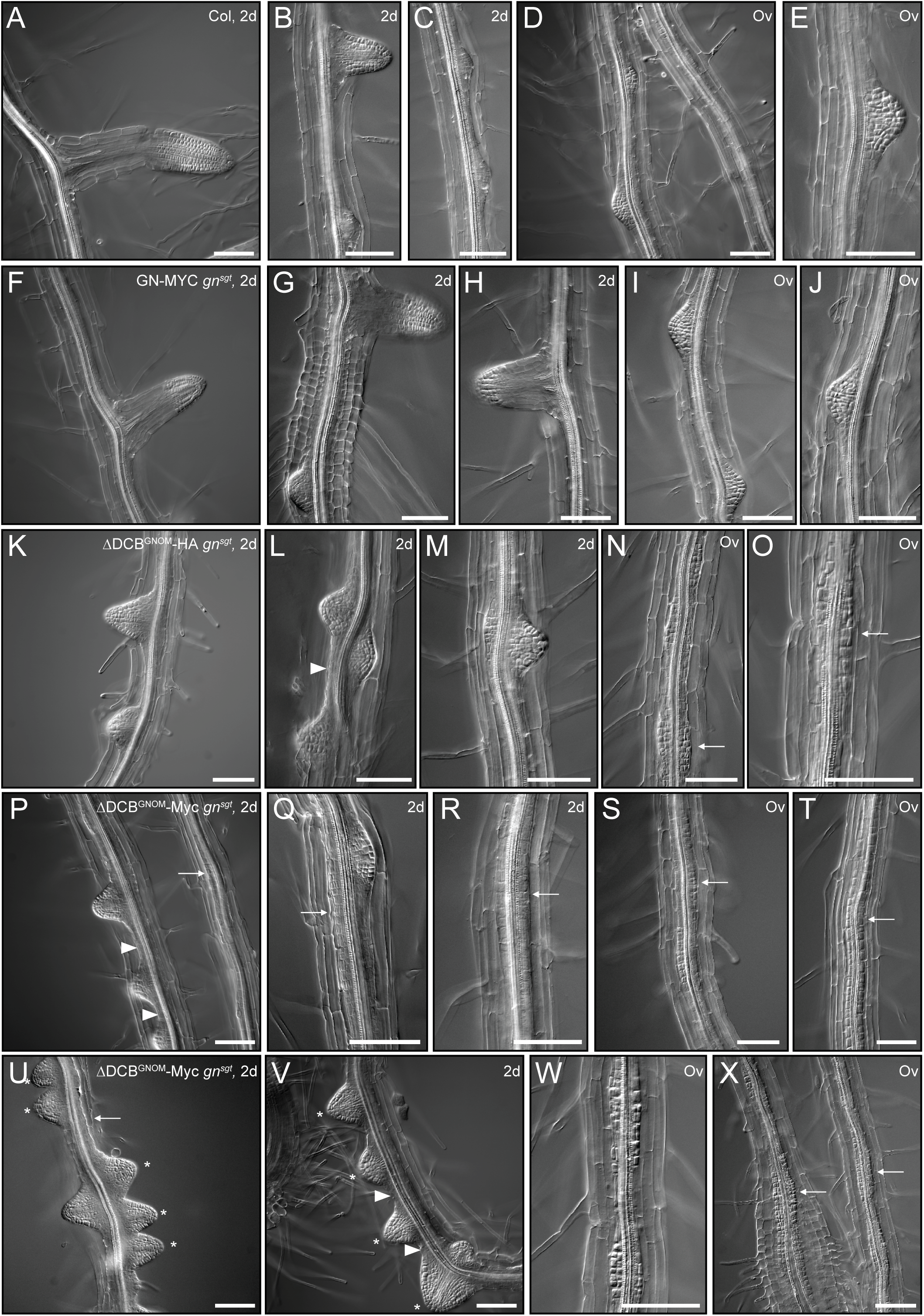
NAA-induced lateral root initiation in *gnom^sgt^* seedlings rescued by ΔDCB^GNOM^. (**A-J**) In Columbia wildtype (Col; **A-E**) and GNOM-Myc *gnom^sgt^* (**F-J**) controls, lateral root primordia formed after NAA treatment for 2 days (2d; **A-C, F-H**) or overnight (Ov; **D-E, I-J**). (**K-X**) In contrast, loss of the DCB domain in GNOM (ΔDCB^GNOM^-HA *gnom^sgt^*, **K-O**; ΔDCB^GNOM^-Myc *gnom^sgt^*, **P-X**) disturbed lateral root formation. After NAA treatment for 2 days (2d; **K-M, P-R, U-V**), some lateral root primordia formed, but were abnormally closely spaced (asterisks) and often pericycle cells strongly proliferated between primordia (arrowhead) or along the whole root axis (arrows). Overnight treatment with NAA (Ov; **N-O, S-T, W-X**) led to strong proliferation of pericycle cells while primordia were rarely detectable (arrows). Scale bar, 100µm.

**Suppl. Table 1.** Rescue of *gnom gnl1* double mutants by *GNOM* transgenes. F1 seedling progeny from reciprocal crosses of *sgt/sgt gnl1/GNL1* plants bearing the transgenes indicated with wild-type (Col) plants were genotyped for the *gnl1* T-DNA allele conferring hygromycin resistance (Hyg^R^) or the hygromycin-sensitive (Hyg^S^) *GNL1* wild-type allele by seed germination of hygromycin-containing agar plates. Myc and HA, protein tags detectable with specific antibodies. ^a^ N, number of seedlings genotyped by PCR ^b^ *sgt* (*gnom^sgt^*), 37-kb deletion spanning *GNOM* and 4 flanking genes on either side (Brumm et al., 2020) ^c^ Transmission of *gnl1* T-DNA allele through pollen reduced by about 40% (Richter et al., 2007) ^d^ Transmission of mutant allele divided by transmission of wild-type allele ^e^ GNOM-GFP protein accumulation at least 10-fold above endogenous GNOM level

| **(A) Parental cross**<br>***Col* (female) X *sgt/sgt gnl1/GNL1* with transgene indicated (male)** | **F1 seedling progeny with paternal alleles (%)** |  |  | **Viability score<sup>d</sup>** |
| --- | --- | --- | --- | --- |
|  | **N<sup>a</sup>** | ***sgt<sup>b</sup> GNL1* (Hyg<sup>S</sup>)** | ***sgt<sup>b</sup> gnl1<sup>c</sup>* (Hyg<sup>R</sup>)** |  |
| *DCB<sup>GNL1</sup>ΔDCB<sup>GNOM</sup>-Myc* (homozygous) | 369 | 63 | 37 | 0.59 |
| *GNOM-Myc* (homozygous) | 307 | 60 | 40 | 0.67 |
| *GNOM<sup>3CS</sup>-YFP* (homozygous) | 203 | 72 | 28 | 0.39 |
| *GNOM-GFP* (homozygous)<sup>e</sup> | 542 | 53 | 47 | 0.90 |
| (values expected for complete rescue) |  | 50 | 50 | 1.0 |
| (values expected for no rescue) |  | 100 | 0 | 0 |

**Suppl. Table 1. Rescue of *gnom gnl1* double mutants by *GNOM* transgenes**
| **(B) Reciprocal parental cross**<br>***sgt/sgt gnl1/GNL1* with transgene indicated (female) X *Col* (male)** | **F1 seedling progeny with maternal alleles (%)** |  |  | **Viability score<sup>d</sup>** |
| --- | --- | --- | --- | --- |
|  | **N<sup>a</sup>** | ***sgt GNL1* (Hyg<sup>S</sup>)** | ***sgt gnl1* (Hyg<sup>R</sup>)** |  |
| *DCB<sup>GNL1</sup>ΔDCB<sup>GNOM</sup>-Myc* (homozygous) | 491 | 51 | 49 | 0.96 |
| *GNOM-Myc* (homozygous) | 274 | 52 | 48 | 0.92 |
| *GNOM<sup>3CS</sup>-YFP* (homozygous) | 228 | 51 | 49 | 0.96 |
| *GNOM-GFP* (homozygous)<sup>e</sup> | 467 | 51 | 49 | 0.96 |
| (values expected for complete rescue) |  | 50 | 50 | 1.0 |

**Suppl. Table 2.** Rescue analysis of *ΔDCB^GNOM^* transgene variants. **(A)** Analysis of *ΔDCB^GNOM^-HA sgt gnl1* mutant gametophyte viability by crossing wild-type (Col) plants with pollen hemizygous for the transgene in the segregating *gnom^sgt^ gnl1* double mutant background. By PCR analysis, no mutant seedlings were doubly heterozygous for *gnom^sgt^* and *gnl1*. HA, protein tag detectable with specific antibody. ^a^ N, number of seedlings genotyped by PCR ^b^ *gnom^sgt^* (*sgt*), 37-kb deletion spanning *GNOM* and 4 flanking genes on either side (Brumm et al., 2020) ^c^ Transmission of *gnl1* T-DNA allele through pollen reduced by about 40% (Richter et al., 2007) ^d^ Assuming independent segregation **(B)** PPT-resistant seedling progeny were analysed for wild-type and *gnom* mutant phenotypes on selection plates (PPT, phosphinotricine). *B4049*, G579R substitution interfering with DCB-ΔDCB interaction and membrane association of GNOM; XLIM, artificial dimerisation module from Xenopus (Anders et al., 2008); Myc, protein tag detectable with specific antibody.

| Parental cross | F1 seedling progeny with paternal alleles (%) |  |  |  |  |
| --- | --- | --- | --- | --- | --- |
|  | N<sup>a</sup> | *GNOM GNL1* | *sgt<sup>b</sup> GNL1* | *GNOM gnl1<sup>c</sup>* | *sgt<sup>b</sup> gnl1<sup>c</sup>* |
| *Col* (female) X $\Delta DCB^{GNOM}$-HA *gnom<sup>sgt</sup>/GNOM gnl1/GNL1* (male) | | | | | |
| $\Delta DCB^{GNOM}$-HA (single copy) | 238 | 38 | 35 | 27 | 0 |
| (values expected for no rescue)<sup>d</sup> |  | 33 | 33 | 33 | 0 |
| (values expected for rescue)<sup>d</sup> |  | 25 | 25 | 25 | 25 |

**Suppl. Table 2. Rescue analysis of $\Delta DCB^{GNOM}$ transgene variants**
| Parental genotype | Seedling progeny<br>Total (N) | Phenotypes (%) |  |
| --- | --- | --- | --- |
|  |  | wild-type | *gnom* |
| *XLIM-ΔDCB<sup>GNOM-B4049</sup>-Myc (PPT-res) gnom<sup>sgt</sup>/GNOM* | 529 | 80 | 20 |
| (values expected for no rescue)<sup>d</sup> |  | 75 | 25 |
| (values expected for rescue)<sup>d</sup> |  | 100 | 0 |

## DATA AVAILABILITY

This manuscript contains Source Data files relating to Figure 1

Figure 1 – Figure Supplement 1

Figure 2

Figure 2 – Figure Supplement 1

Figure 2 – Figure Supplement 4

Figure 3

Figure 4

